# Actin-Driven Nanotopography Promotes Stable Integrin Adhesion Formation in Developing Tissue

**DOI:** 10.1101/2023.07.28.550203

**Authors:** Tianchi Chen, Cecilia Huertas Fernández-Espartero, Abigail Illand, Ching-Ting Tsai, Yang Yang, Benjamin Klapholz, Pierre Jouchet, Mélanie Fabre, Olivier Rossier, Bianxiao Cui, Sandrine Lévêque-Fort, Nicholas H. Brown, Grégory Giannone

## Abstract

Morphogenesis requires building stable macromolecular structures from highly dynamic proteins. Muscles are anchored by long-lasting integrin adhesions to resist contractile force. However, the mechanisms governing integrin diffusion, immobilization, and activation within developing tissues remain elusive. Here, we show that actin polymerisation-driven membrane protrusions form nanotopographies that enable strong adhesion at Drosophila muscle attachment sites (MAS). Super-resolution microscopy revealed that integrins assemble adhesive belts around Arp2/3-dependent actin protrusions, forming invadosome-like structures with membrane nanotopographies. Single protein tracking showed that, during MAS development, integrins became immobile and confined within diffusion traps formed by the membrane nanotopographies. Actin filaments also displayed restricted motion and confinement, indicating strong mechanical connection with integrins. Using isolated muscles cells, we show that substrate nanotopography, rather than rigidity, drives adhesion maturation by regulating actin protrusion, integrin diffusion and immobilization. These results thus demonstrate that actin-polymerisation driven membrane protrusions are essential for the formation of strong integrin adhesions sites in the developing embryo, and highlight the important contribution of geometry to morphogenesis.

## Introduction

A fundamental question in biology is how to build stable macromolecular structures from highly dynamic proteins. Cell-cell and cell-matrix adhesions are such structures important for cell migration during tissue morphogenesis, but are also crucial for their mechanical stability ^1,2^. In particular, muscles must be connected by long-lasting adhesive structures to tendons, tendon cells, or other muscle cells to resist the force of muscle contraction ^3–5^. Integrin adhesive structures are composed of a diverse but evolutionarily conserved set of proteins that link the actin cytoskeleton to the extracellular matrix through transmembrane integrin dimers ^6–8^. Actin motion and myosin contractility are able to generate force on the integrin adhesions through molecular mechanosensors such as talin ^9^ and vinculin ^10^, forming a molecular clutch where actin flow speed and clutch strength are anti-correlated ^11–14^. The nanoscale organization and molecular dynamics of integrin adhesions have been well studied, especially on flat non-deformable substrates ^15–19^. However, cells *in vivo* reside in a three-dimensional (3D), dynamic, deformable, and confined tissue environment resulting in different molecular organization and biomechanical control of adhesion formation ^20–23^. Here we focus on the formation of Drosophila muscle attachment sites (MAS), which are the most prominent integrin adhesive structures in the developing embryo ^4^. They connect the muscles to specialized epidermal cells, the tendon cells, to anchor their contraction to the apical extracellular matrix. This *in vivo* model differs from integrin adhesions formed by cells in culture in fundamental ways. Electron microscopy showed that MASs are not flat structures, but instead are extensively interdigitated ^24^, which could increase the strength of the attachment by increasing the surface area of the contact. In addition, membrane molecular organization and nanotopographies can impact the dynamics and clustering of membrane proteins, including integrins ^25–28^. Although mechanical force is clearly important for the formation of a fully functional MAS ^29–31;^ whether actin polymerisation provides that force to an integrin molecular clutch, as is essential in the formation of integrin adhesions on flat surfaces in culture, is not known. Thus, it remains unclear at the molecular level in a complex 3D tissue environment, how membrane and actin network dynamics regulate the diffusion, immobilization and activation of integrins to trigger the formation of mechanically stable and mature adhesions.

To address these questions, we used an organ culture model of MAS formation compatible with super-resolution microscopy and single protein tracking. We found that integrins assemble adhesive belts around Arp2/3-dependent actin protrusions in an invadosome-like structure with nanotopographical patterns. Inhibition of actin protrusions abolished MAS development and disassembled mature MASs. MAS maturation was also accompanied by decreased actin filament movement, whereas integrin displayed increased molecular confinement and immobilization. We found these changes to integrin behavior were the result of diffusion traps generated by 3D nanotopography induced by invadosome-like actin protrusions. Using primary culture of muscle cells, we show that substrate nanotopography, rather than substrate stiffness, promotes the formation of stable MASs, by regulating integrin diffusion and immobilization. These results emphasize the importance of actin-protrusion generated nanotopography in promoting the formation of stable integrin adhesions in developing tissue, and highlight the essential role of geometrical information for morphogenesis.

## Results

### Super-resolution microscopy reveals 3D invadosome-like adhesion structure

Examination of the formation of MASs has been hindered by the thickness of the embryo, preventing high spatial and temporal resolution imaging, the morphogenetic movements of the embryos, preventing long-term tracking of developing MASs, and the vitelline membrane surrounding the embryo, which prevents the straightforward use of inhibitors. Here, we circumvented these problems by dissecting embryos to produce *ex vivo* embryonic fillets ^32^ that can be cultured for the few hours of MAS development. We dissected early stage 16 embryos in culture medium, and attached the apical surface of the epidermis to the coverslip (**Fig. 1a**, also See **Methods** & **Extended Data Fig. 1a-d**). The main advantage of this is the proximity to the coverslip and the immobilisation of the tissue until the muscles become highly contractile. Importantly, the MASs continue their maturation (**Fig. 1a, b** and **Supplementary Movie 1, 2**) and the muscles develop spontaneous contractions normally (**Supplementary Movie 3, 4**). We focused on the MASs formed by the lateral-transverse body wall muscles (LT1-3), which are disc-shaped adhesions parallel to the coverslip, and about 5-10 µm away (**Fig. 1a** and **Extended Data Fig. 1b, c**). Time-lapse imaging of developing embryo fillets showed nascent adhesions with isolated dot-like morphology, which then transformed into mature adhesions with enrichment of integrin (βPS-GFP) and paxillin (paxillin-GFP) (**Extended Data Fig. 1e-f** and **Supplementary Movie 2**), within an area of 17.9±0.6 µm^2^ (mean ± s.e.m., 4.5h *ex vivo*) (**Extended Data Fig. 1g**).

**Figure 1.**
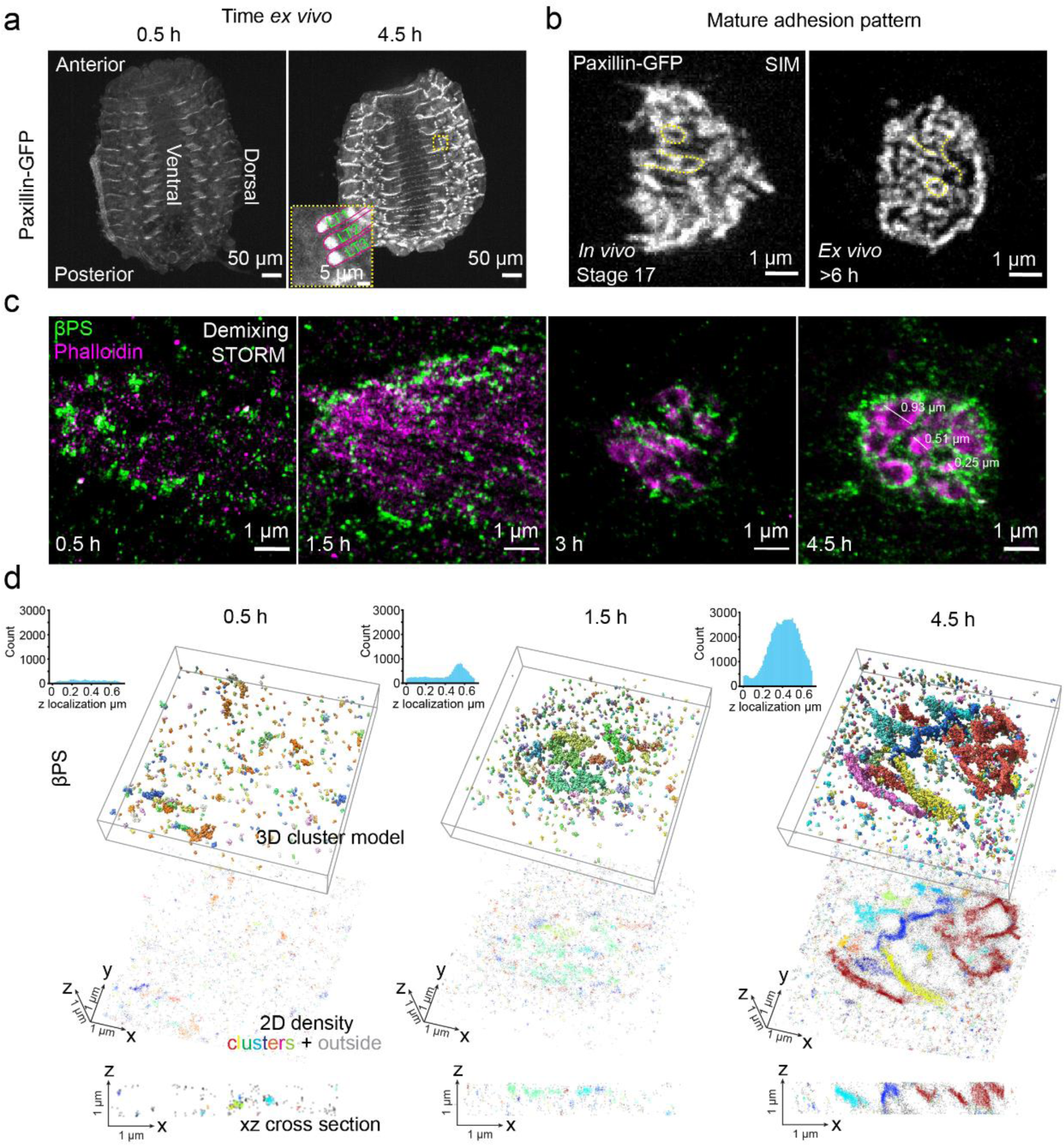
Actin protrusion induces invadosome-like integrin adhesion and pattern formation in 3D tissue development of Drosophila muscle attachment site. **a,** *Ex vivo* embryo fillet culture preserves muscle attachment site (MAS) development and enrichment of the adhesion protein Paxillin-GFP. Inset shows a set of MASs from the lateral-transverse muscles (LT1-3). Scale bars, 50 μm. **b,** SIM imaging shows similarity between finger-print-like patterns in a mature MAS *in vivo* (left) and *ex vivo* (right). Scale bars, 1 μm. **c,** Development of the finger-print-like pattern revealed by two-color-demixing STORM images of actin (magenta) and integrin (green) clusters. Scale bars, 1 μm. **d,** Adhesions increase in depth, as revealed by 3D ModLoc super-resolution imaging of integrin clusters in developing MASs, presented as surface models (top), 2D projections (centre) and xz cross sections (bottom), with each cluster coloured randomly and signal outside the clusters coloured grey. Histograms show z-distribution of integrins.

The cultured embryo fillets allowed us to perform quantitative super-resolution microscopy and single protein tracking, and to use pharmacological treatments that target the acto-myosin cytoskeleton and integrins. 2D and 3D structured illumination microscopy (SIM) imaging of mature adhesion sites in late-stage embryo fillets, and in intact stage 17 embryos, revealed that integrins are patterned into ridge-like structures, resembling a fingerprint (**Fig. 1b**, **Extended Data Fig. 1h**, and **Extended Data Fig. 2a-b**). Two-colour super-resolution imaging with spectral demixing STORM further unveiled the nanoscale organization of MASs and actin. Early MASs exhibit isolated clusters of integrin that, as the MASs mature, become organized into ridge-shaped structures, with integrin clusters surrounding an actin core (**Fig. 1c** and **Extended Data Fig. 2a-b**). These are reminiscent of invadosome-like structures ^21,33^. With ModLoc 3D STORM super-resolution imaging ^34^ of different stages of MAS formation, we also observed the transition of integrins from isolated clusters to more elongated structures within the convoluted membrane that extend in the z direction (**Fig. 1d**, **Extended Data Fig. 2c** and **Supplementary Movie 5**). These findings prompted us to examine the role of actin in the formation of this invadosome-like structure, and whether integrin adhesion proteins display altered mobility and stability within the ridges.

### Arp2/3-dependent actin protrusions drive adhesion development in MAS

Mechanical forces generated by acto-myosin contraction and actin flow are essential for integrin adhesion initiation and maturation in cells in culture ^14,35–37^. In the MAS, although muscle and non-muscle myosin generate forces that regulate adhesion reinforcement and protein turn-over ^29,31^, neither is required for the initial development of the adhesion before muscle contraction ^31,38,39^. Consistent with these findings, when we inhibited Rho-kinase activity with Y-27632 in the embryo fillet, the amount of integrin was reduced but the MAS still matured and formed fingerprint patterns (**Extended Data Fig. 3a, b**). This shows that non-muscle myosin contractility is not essential for MAS development but contributes to its force-dependent reinforcement.

We then tested whether the force from actin polymerization is important in this system, which has not been addressed with any genetic or imaging approaches. In cells in culture, the connection of integrins to the retrograde actin flow in cell protrusions, such as the lamellipodium, initiates early adhesions ^13,40^. This flow is driven by the polymerization of branched actin networks, initiated by the Arp2/3 complex (Arp2/3) ^41,42^. We found that inhibition of Arp2/3 by continuous CK666 treatment completely abolished MAS maturation (**Extended Data Fig. 3c, d**), and disassembled mature MASs (**Fig. 2a, b** and **Supplementary Movie. 6**). Live imaging (every 10 min) throughout MAS formation revealed small actin protrusions in nascent adhesions, which transformed into more pronounced and stable protrusions in mature adhesions (**Fig. 2c-e**, **Extended Data Fig. 3e**, and **Supplementary Movie. 7**). These actin protrusions may trigger integrin movement and reorganization observed during MAS formation (**Extended Data Fig. 3f** and **Supplementary Movie 8**). The growth and pattern formation of the MAS are thus driven by actin protrusions that recruit integrin around them (**Fig. 2f**).

**Figure 2.**
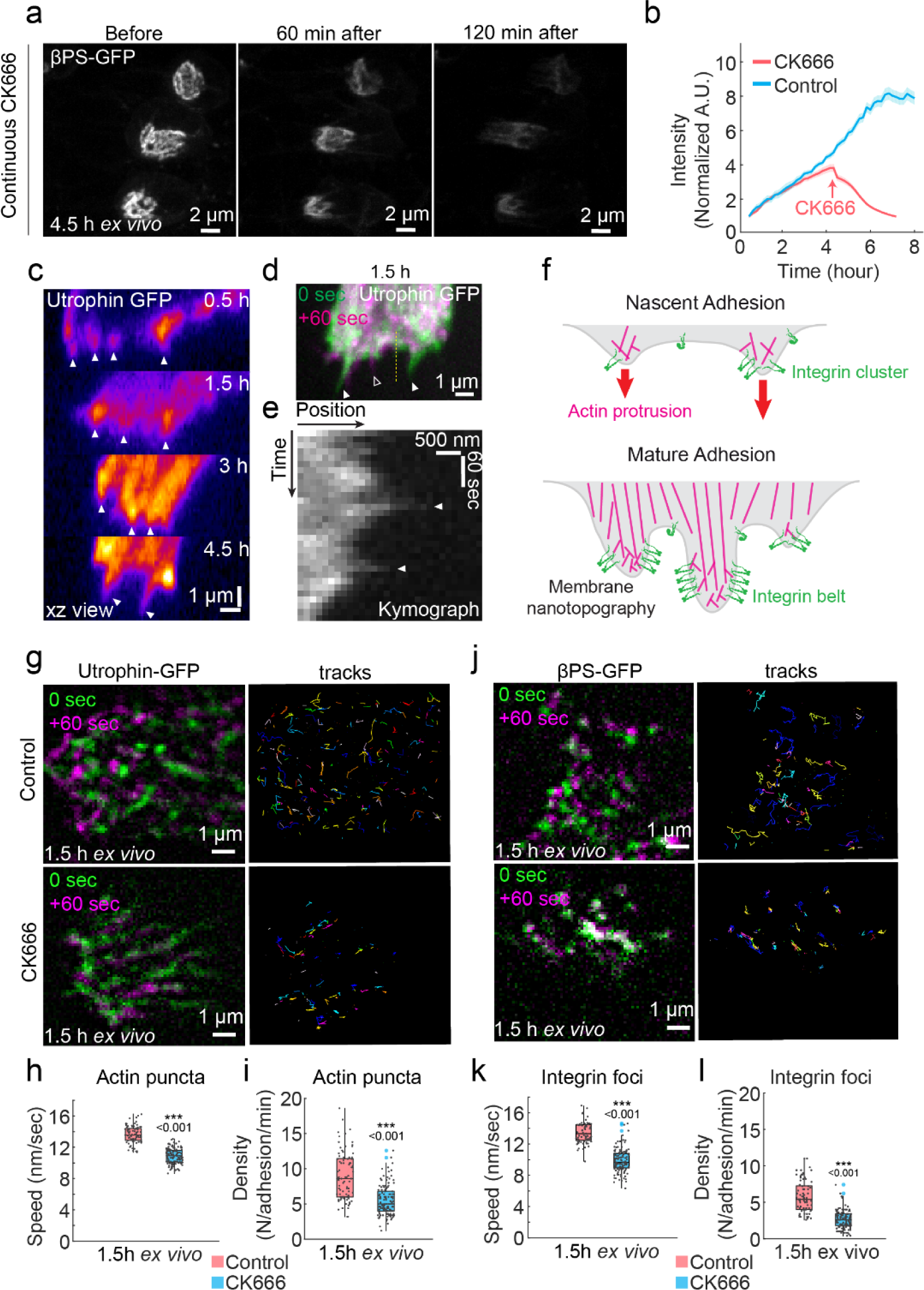
Adhesion development and integrin dynamics driven by Arp2/3-dependent actin polymerization. **a,** Continuous treatment of CK666 in mature adhesions (4.5h *ex vivo*) induces their disassembly. Scale bars, 2 μm. b, Quantification of ꞵPS-GFP fluorescence intensity of control and CK666 treated adhesions (mean ± s.e.m.). **c**, xz view of actin protrusions (filled arrow head) in a developing MAS, actin labelled by expression of the actin-binding domain of Utrophin tagged with GFP, Horizontal scale bar, 1 μm. Vertical scale bar, 1 μm. **d**, Overlay showing dynamic actin protrusions 60 seconds apart, from a 1.5h MAS *ex vivo*. Solid arrowhead shows protrusions at 0 sec in green; outlined arrowhead shows a protrusion at +60 sec in magenta. Scale bar, 1 μm. **e**, Kymograph along yellow dashed line in d. Horizontal scale bar, 500 nm. Vertical scale bar, 60 sec. **f**, Schematic of the organization of actin (magenta) and integrin (green) in nascent and mature adhesions. **g**, Utrophin-GFP labelled actin puncta and puncta tracks in control and CK666 treated adhesion (1.5h *ex vivo*). Scale bars, 1 μm. **h**, Movement speed of actin puncta. **i**, Density of actin puncta. **j**, Integrin foci and tracks (βPS-GFP) in control and CK666 treated MASs (1.5h *ex vivo*). Scale bars, 1 μm. **k**, Movement speed of integrin foci. **l**, Density of integrin foci.

Using higher temporal resolution imaging (0.1 Hz) at different stages of MAS formation, we also observed highly dynamic actin puncta and integrin foci in nascent adhesions (1.5 h *ex vivo*) (**Fig. 2g, j**) and the formation of more stable structures in mature adhesions (4.5h *ex vivo*) (**Extended Data Fig. 4a, b** and **Supplementary Movie 9**). The speed of both actin puncta and integrin foci decreased during MAS development (**Extended Data Fig. 4c, d**), accompanied by a transition from highly dynamic protrusions to less dense but more stable protrusive structures, lasting 27.7±0.3 seconds at 1.5h to 49.8±1.0 seconds at 4.5h (mean ± s.e.m.) (**Extended Data Fig. 4e, f**). The actin puncta are formed by bursts of actin polymerization, since their speed and density were reduced by CK666 treatment (**Fig. 2g-i** and **Supplementary Movie 10**). Consistently, expression of the Arp2/3 complex component Arp3-GFP, or a component of the WAVE regulatory complex, WAVE-eGFP ^43,44^, in the muscles, showed similar dynamic puncta within developing adhesions (**Extended Data Fig. 4g-i**). CK666 treatment also reduced the speed and density of integrin foci (**Fig. 2j-l** and **Supplementary Movie 11**). The coordinated assembly and movement of Arp2/3-dependent actin puncta and integrin foci suggest mechanical coupling between the two and the reduction of movement speed during adhesion development indicates the enhancement of their molecular connection^13,45^.

### Developing MAS regulates integrin adhesion protein diffusion and immobilization

Membrane nanotopography generated by the actin cytoskeleton and large membrane glycoproteins can affect membrane protein diffusion and clustering ^25,27,46^. We thus investigated whether the formation of the fingerprint ridges alters integrin diffusion. We performed high-frequency (50 Hz) single protein tracking coupled with photo-activation localization microscopy (sptPALM), at 1.5h, 3h, and 4.5h (**Supplementary Movie 12,** see Methods for detail). We used mean square displacement (MSD) analysis to quantify protein diffusive properties including diffusion mode and diffusion coefficient (D) ^7,17,18^ (**Fig. 3a-e** and **Extended Data Fig. 5a-e**). As observed in mature integrin adhesions in cultured cells ^17,18,47^, a significant fraction of integrins (ꞵPS-mEos3.2) remained diffusive inside the adhesion during its development, and the fraction decreased with time (**Fig. 3a, d**; 73.7±1.6% at 1.5h, 49.0±1.3% at 4.5h, mean ± s.e.m.). Thus, integrin immobilization increased with MAS maturation in contrast to membrane targeted mEos3.2-CAAX, which continued to diffuse freely inside the MAS with minimal immobilization (**Fig. 3b, d**, **Extended Data Fig. 5i, j**, and **Extended Data Fig. 6a-d**). Integrin diffusion coefficient decreased during adhesion maturation (0.117±0.003µm^2^/sec at 1.5h, 0.084±0.004µm^2^/sec at 4.5h, mean ± s.e.m.) (**Fig. 3e** and **Extended Data Fig. 5d, e**). The diffusion coefficient of mEos3.2-CAAX, though higher than that of integrin, also decreased during adhesion development (0.383±0.011µm^2^/sec at 1.5h, 0.265±0.006µm^2^/sec at 4.5h, mean ± s.e.m.) (**Fig. 3e** and **Extended Data Fig. 6c, d**). These results suggest that the increased efficiency of integrin immobilization in MAS is correlated with the formation of the fingerprint ridges and change in diffusion dynamics.

**Figure 3.**
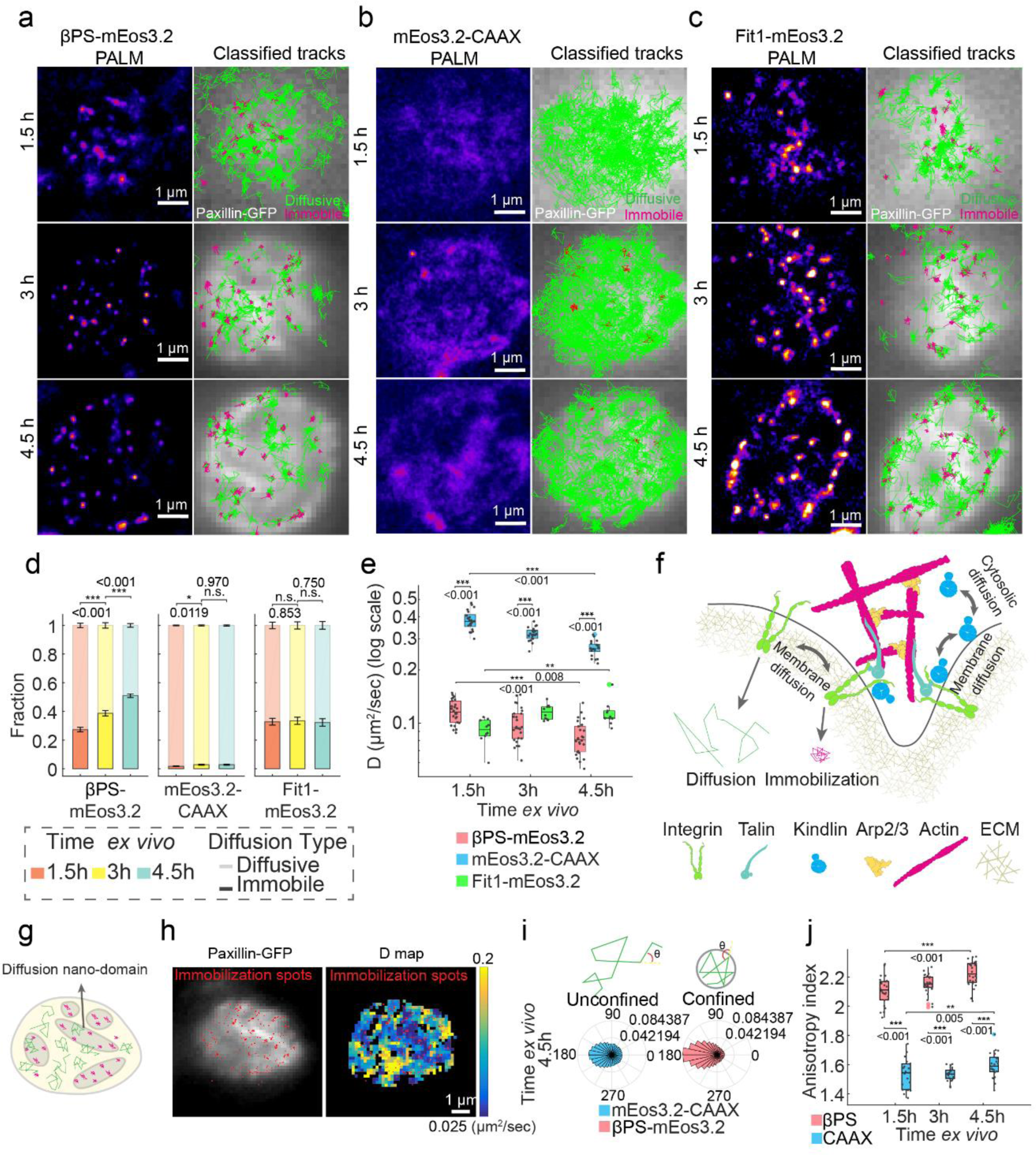
Diffusion and confinement of integrin adhesion protein dynamics during MAS development. **a-c**, sptPALM tracking of integrin adhesion molecules and membrane anchored CAAX reveal their diffusion trajectories inside developing MASs. Left columns, reconstructed PALM images of ꞵPS-mEos3.2, mEos3.2-CAAX and Fit1-mEos3.2 molecular localizations. Right columns, individual tracks are classified into diffusive (green) or immobile (magenta). Scale bars, 1 μm. **d**, Fraction of immobile and diffusive trajectories for ꞵPS-mEos3.2, mEos3.2-CAAX and Fit1-mEos3.2 (mean ± s.e.m.). **e**, Diffusion coefficient (D) of diffusive ꞵPS-mEos3.2, mEos3.2-CAAX and Fit1-mEos3.2 molecules. **f**, Schematic showing adhesion protein diffusion and immobilization in developing MAS. Whereas integrin diffusion is confined on the membrane, Kindlin can exchange between membrane and cytosolic diffusion. **g**, Schematic of the organization of diffusion nano-domains in mature adhesions with diffusing (green) and immobilized (magenta) tracks of integrin molecules. **h**, Representative image showing the correlation between Paxillin-GFP image (left) and diffusion coefficient map (right). Immobilization spots are represented as red dots. Scale bar, 1 μm. **i**, Distribution of angles between steps in the tracks for CAAX-mEos3.2 (left column) and ꞵPS-mEos3.2 (right column). Schematic illustrates freely diffusing trajectory (left) vs. spatially confined trajectory (right). **j**, Anisotropy of step angles for CAAX-mEos3.2 and ꞵPS-mEos3.2.

Importantly, the integrin activator kindlin (Fermitin1/Fit1 in *Drosophila*) exhibited different diffusive behaviour inside developing MAS **(Fig. 3c** and **Extended Data Fig. 7a-e**). Whereas ꞵPS-mEos3.2 showed increased immobilization, Fit1-mEos3.2 maintained a steady level of immobilization during adhesion development (**Fig. 3d**). Furthermore, whereas the two membrane inserted molecules, ꞵPS-mEos3.2 and mEos3.2-CAAX, showed decreased diffusion coefficient and increased confinement, this was not the case for Fit1-mEos3.2 which showed an increase of diffusion coefficient (**Fig. 3e**). This suggests the existence of diffusion barriers in the developing MAS restricting the movement of membrane proteins, while the transient interactions of kindlin with the membrane, via its PH domain binding to PIP2 ^18^, allows a more dynamic switch between cytosolic and membrane diffusion (**Fig. 3f**).

### Diffusion nano-domains confine adhesion protein movement

The organization of adhesion into ridge-like structures and restricted diffusion of integrin suggest the formation of diffusion nano-domains inside MAS (**Fig. 3g**). Indeed, within mature adhesions, we observed distinct zones with diffusing integrins displaying lower diffusion coefficient and step size, which corresponded to regions of high paxillin-GFP intensity and increased integrin immobilization (**Fig. 3h** and **Extended Data Fig. 5f**). Thus, the MAS is segregated into nano-domains of rapid versus restricted diffusion. To quantify the confinement of integrin, we calculated the step angle anisotropy ^48^ (see Methods) of diffusing ꞵPS-mEos3.2 tracks, which exhibited highly anisotropic distribution of step angles, compared to mEos3.2-CAAX, indicating a confined behaviour (**Fig. 3i-j**, **Extended Data Fig. 5g, h** and **Extended Data Fig. 6e, f**). The step angle anisotropy of both ꞵPS-mEos3.2 and mEos3.2-CAAX increased during MAS development (**Fig. 3j**), suggesting increasing geometrical confinement. The anisotropy index increased with larger step size (>400 nm), comparable to the size of the ridges (**Fig. 1c**), compared to smaller step size (100 nm) for both ꞵPS-mEos3.2 and mEos3.2-CAAX (**Extended Data Fig. 5h** and **Extended Data Fig. 6f**). This suggests that the effect of confinement is due to the geometry of membrane nano-domains within MAS. In contrast, kindlin experiences similar levels of confinement during MAS maturation (**Extended Data Fig. 7e**), suggesting it can bypass membrane geometrical confinement with cytosolic diffusion (**Fig. 3f**). In the cell body, where the protrusive structures are not present, integrins diffuse faster (0.250±0.003 µm^2^/sec, mean ± s.e.m.) with decreased immobilization fraction (27.3±1.6 %, mean ± s.e.m.) and reduced confinement compared to inside MAS (**Extended Data Fig. 8a-j**), suggesting that the restricted diffusion dynamics of integrin is specific to the MAS. Altogether, these results indicate that actin protrusions and the formation of ridge-like structures during MAS maturation could serve as diffusion traps that concentrate membrane proteins and promote their interaction, immobilization and clustering (**Fig. 3f**) ^49–51^.

### Confined actin movement in MAS forms a strong connection with integrin adhesion

The increased immobilization of integrin during adhesion maturation also suggests stronger engagement with the actin cytoskeleton ^17,45,52^. We thus investigated actin filaments (F-actin) slow movement with low frequency (2 Hz) sptPALM ^17,53^ of actin-mEos3.2 inside MAS (**Fig. 4a** and **Extended Data Fig. 9a**). We first distinguished moving and stationary F-actin and observed an increase of the stationary fraction during MAS development (**Fig. 4b**). In addition, the moving fraction displayed isotropic distribution of angles (**Fig. 4a** and **Extended Data Fig. 9b**), with mainly highly confined movements, other than directed or Brownian motion (**Fig. 4c-d**). Furthermore, the speed of F-actin movement decreased during MAS development (**Fig. 4e**). CK666 treatment also led to a reduction of F-actin movement (**Fig. 4e** and **Extended Data Fig. 9c-d**). These results are in contrast to the coordinated retrograde F-actin flow observed in the lamellipodium, site of adhesion initiation, and mature integrin adhesions of migrating cells ^13,17,40,54,55^. To probe the activation status of integrin during MAS development, we performed Mn^2+^ stimulation to activate integrins ^17,56,57^ at different stages (**Extended Data Fig. 10a-c**) and observed increased immobile fraction in nascent adhesions but not in mature ones (**Fig. 4f** and **Extended Data Fig. 10d**), suggesting increased level of integrin activation during MAS development. As expected for diffusing integrins, only minor effects were seen on the diffusion coefficient and confinement after Mn^2+^ stimulation (**Extended Fig. 10e-g**). The increasing fraction of stationary F-actin and decreasing rate of F-actin movement during MAS development, together with increased integrin immobilization and activation, is consistent with the formation of a stable molecular connection during adhesion maturation (**Fig. 4g**).

**Figure 4.**
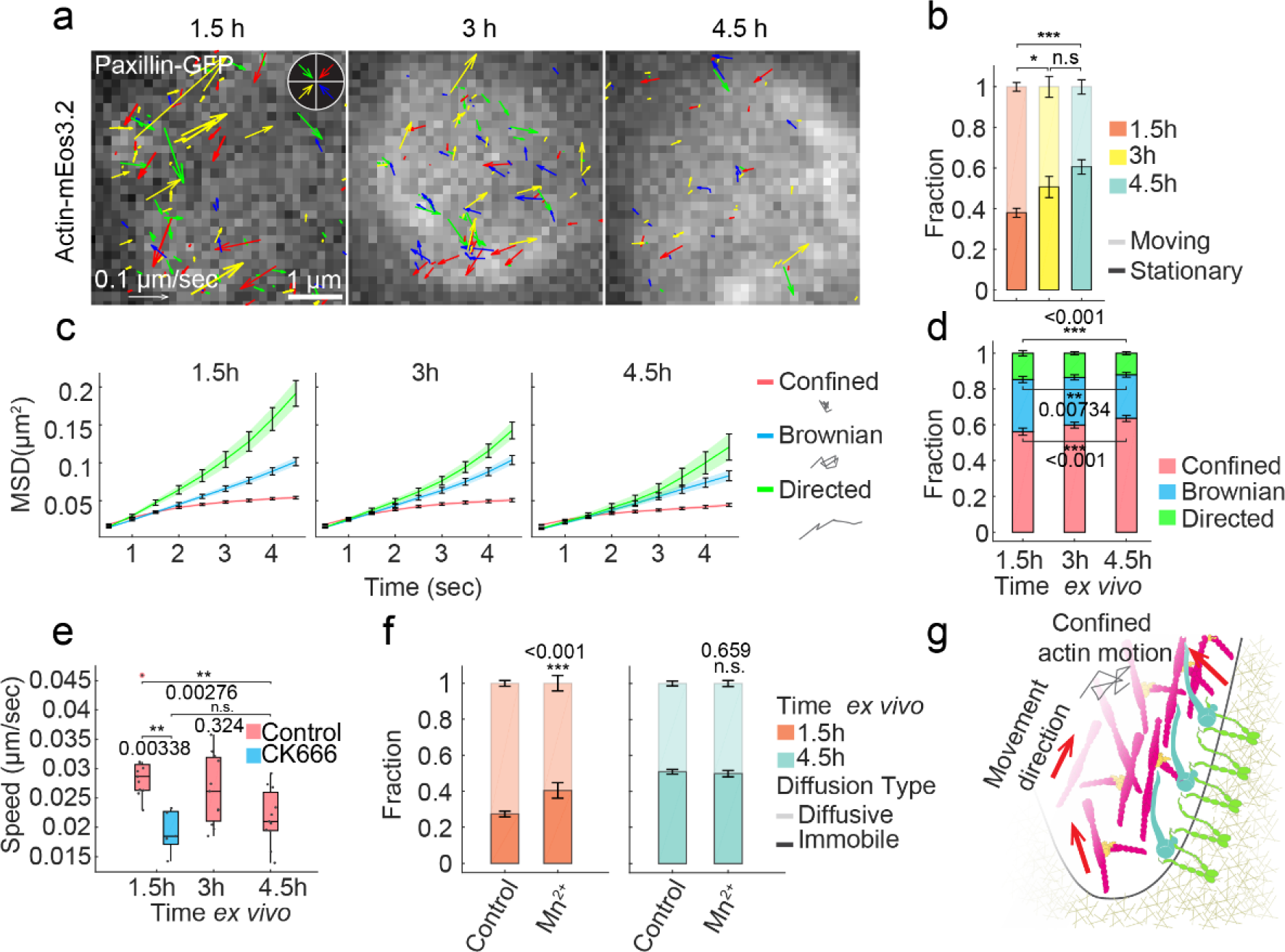
Confined actin motion promotes strong molecular engagement with integrin adhesion. **a**, F-actin movement measured from tracking of Actin-mEos3.2 shows isotropic distribution of direction in developing MASs, rather than a coordinated flow from the muscle end. Scale bars, left: 0.1 μm/sec, right: 1 μm. **b**, Fraction of stationary and moving actin-mEos3.2 tracks quantifies reduction in actin polymerization. n = 8 embryos for 1.5h, 3h and 4.5h. **c**, Mean square displacement curves of moving actin-mEos3.2 tracks showing directed, Brownian, or confined motion. **d**, Fraction of moving Actin-mEos3.2 tracks exhibiting directed, Brownian, or confined motion. **e**, Actin movement speed for control at 1.5h, 3h and 4.5h and CK666 treatment at 1.5h. n = 11 embryos for control 1.5h, 3h and 4.5h. n = 5 for CK666 1.5h. **f**, Diffusive and immobile fraction of ꞵPS-mEos3.2 for control and Mn^2+^ stimulation. n = 26 embryos for control 1.5h and 4.5h. n = 9 embryos for Mn^2+^ stimulation 1.5h and 4.5h. **g**, Illustration showing confined actin movement and molecular engagement with integrin adhesion in developing MAS.

### Dynamic actin protrusion and stable actin structure regulate integrin diffusion during MAS development

We then tested whether integrin diffusion and immobilization are controlled by F-actin networks in MAS. Continuous incubation with Latrunculin A or CK666 abolished the maturation of MASs, decreased the immobilization and confinement of integrin, and increased its diffusion coefficient (**Fig. 5a-d**, **Extended Data Fig. 11a-I** and **Extended Data Fig. 12a-f**). This was specific to the MAS, unlike the cell body devoid of actin protrusions where integrin diffusion was unaffected by CK666 treatment (**Extended Data Fig. 13a-h**). Changes in integrin diffusive behaviour triggered by Arp2/3 inhibition were only partially rescued by Mn^2+^ stimulation (**Fig. 5b-d** and **Extended Data Fig. 12b-f**), indicating that direct actin-dependent integrin activation cannot solely account for the effects on integrin diffusion regulation.

**Figure 5.**
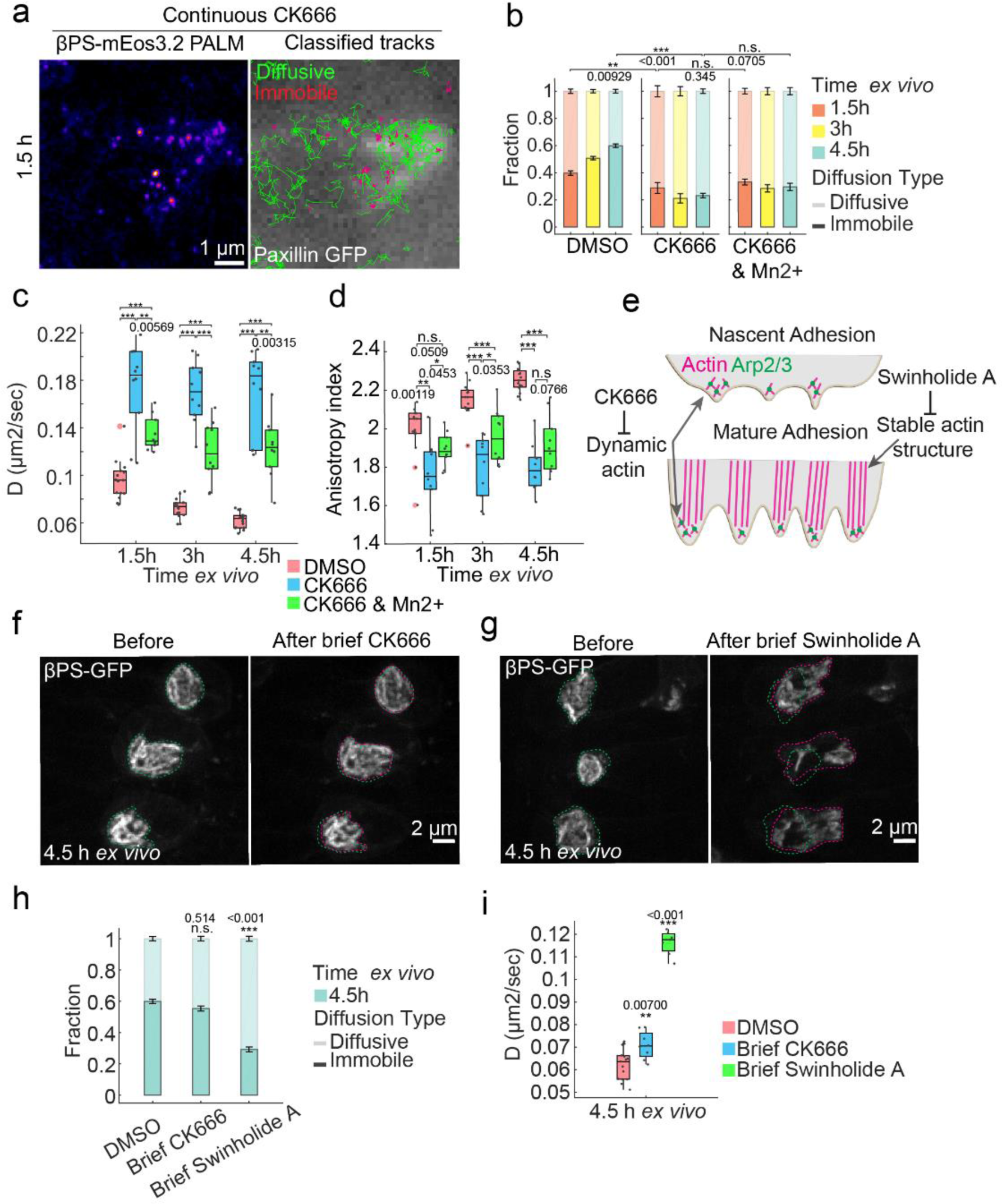
Actin dynamics and stable structure regulate integrin diffusion dynamics during adhesion maturation. **a**, sptPALM tracking of integrin (ꞵPS-mEos3.2) with continuous CK666 treatment. Scale bar, 1 μm. **b**, Fraction of immobile and diffusive trajectories for ꞵPS-mEos3.2. n = 15 embryos for DMSO, n = 10 for CK666, n = 10 for CK666 & Mn^2+^, same for c, d. **c**, Diffusion coefficient of diffusive ꞵPS-mEos3.2 molecules. **d**, Step angle anisotropy for ꞵPS-mEos3.2. **e**, Schematic showing the effect of CK666 on dynamic actin in nascent and mature adhesions vs. the effect of Swinholide A on stable actin structure in mature adhesions. **f**, Adhesion morphology before (green dashed line) and after (magenta dashed line) brief CK666 treatment in mature adhesions. Scale bar, 2 μm. **g**, Adhesion morphology before (green dashed line) and after (magenta dashed line) brief Swinholide A treatment in mature adhesions. Scale bar, 2 μm. **h**, Fraction of immobile and Diffusive trajectories for ꞵPS-mEos3.2 (mean ± s.e.m.). n = 15 embryos for DMSO, n = 8 for CK666, n = 7 for Swinholide A, same for i. **i**, Diffusion coefficient of diffusive ꞵPS-mEos3.2 molecules.

We thus tested whether F-actin networks form a stable structural scaffold in the MAS that could impose geometric constraint on the membrane and be responsible for the changes in integrin diffusion. We distinguished dynamic from stable actin by performing brief and continuous drug inhibitions in nascent and mature MASs (**Fig. 5e**, **Extended Data Fig. 14a-g** and **Extended Data Fig. 15a-f**). The effect of brief Arp2/3 inhibition on nascent (1.5h) MASs was quantified 15 minutes after CK666 treatment (**Extended Data Fig. 14a**). This reduced the diffusive fraction of integrins, increased their diffusion coefficient and reduced integrin confinement (**Extended Data Fig. 14b-g**). However, when mature adhesions (4.5h) were briefly treated with CK666, MAS morphology was unaffected (**Fig. 5f and Extended Data Fig. 15a**), and the immobile fraction was unaltered (**Fig. 5h**). The contrast between continuous and brief CK666 treatment suggests that Arp2/3-induced actin polymerization is responsible for a dynamic pool of actin filament (**Fig. 2c-f**) both driving dynamic membrane protrusions especially in nascent adhesions and the maintenance of a more stable actin structures in mature adhesions, since continuous inhibition of Arp2/3 in mature adhesions led to their disassembly (**Fig. 2a-b**). On the other hand, when mature MAS were treated with Swinholide A, which severs actin filaments, the adhesion structure was immediately perturbed (**Fig. 5g**, **Extended Data Fig. 15b** and **Movie. 13**), resulting in increased diffusive fraction and diffusion coefficient of integrin (**Fig. 5h, i** and **Extended Data Fig. 15c-e**). Together, these results indicate that the coordination of dynamic actin polymerization and more stable F-actin structures build a 3D scaffold controlling integrin diffusion and immobilization during MAS formation.

### Substrate nanotopography promotes adhesion maturation by regulating actin protrusion and integrin diffusion

Substrate nanotopography and curvature can regulate integrin adhesion and actin dynamics ^25,58,59^. In addition, substrate stiffness is also an important parameter controlling engagement of the integrin molecular clutch and adhesion formation ^60–63^. We thus hypothesized that the restriction of integrin diffusion induced by actin protrusions in the deformable tissue may be a result of its physical properties such as rigidity and topography.

In order to test this hypothesis, we isolated and cultured muscles from mid stage 16 embryo fillets and placed them on substrates with different biophysical properties. On rigid and flat substrates coated with fibronectin, muscles only formed nascent dot-like adhesions that did not mature (**Fig. 6a**). Reducing substrate stiffness did not promote the formation of mature MASs, as isolated muscles only formed nascent adhesions on polydimethylsiloxane (PDMS) gels with stiffnesses of 3 kPa, 15 kPa and 35 kPa (**Extended Data Fig. 16a**). The intensity of ꞵPS-GFP even decreased on softer substrates (**Extended Data Fig. 16b**), consistent with studies of mechanosensitive integrin adhesions in cultured cells ^61–63^. These nascent adhesions were not able to withstand contractile forces, as the muscles detached during spontaneous muscle contractions (**Extended Data Fig. 16c** and **Supplementary Movie. 14**). The reduced adhesion maturation was not due to lower expression of integrin on soft gels, as we observed that adhesions could form when muscles contacted each other (**Extended Data Fig. 16d**). Thus, the rigidity of the muscle-tendon interface is not a determinant factor of MAS maturation and stabilisation.

**Figure 6.**
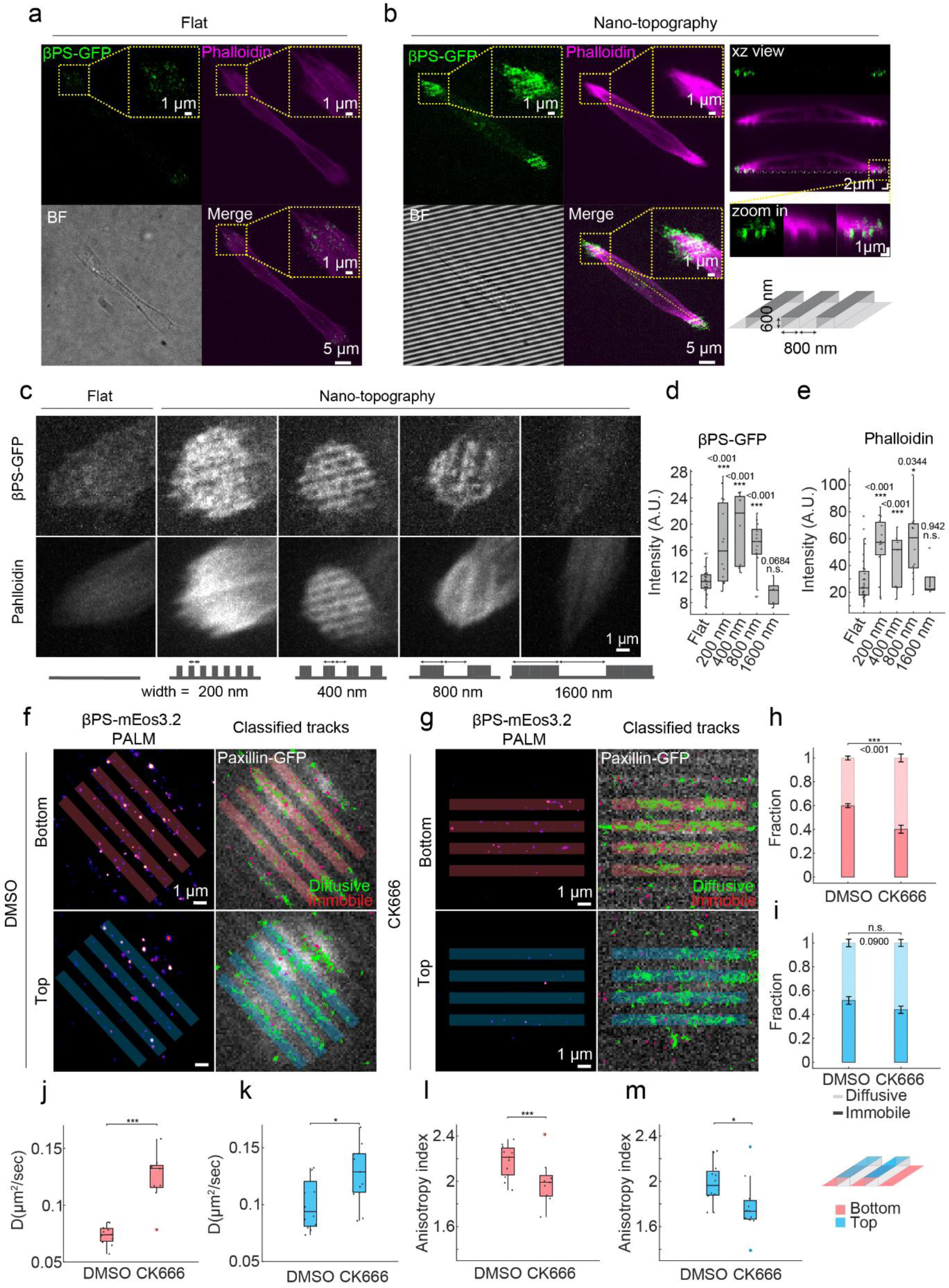
Nanotopography promotes adhesion formation and confinement of integrin diffusion through actin cytoskeleton. **a**, Isolated muscle cell on rigid flat substrate expressing ꞵPS-GFP (green) and stained with Phalloidin (magenta). Scale bar, 5 μm. zoom in panel scale bars, 1 μm **b**, Isolated muscle cell on nanopatterned substrate expressing ꞵPS-GFP (green) and stained with Phalloidin (magenta). Scale bars 5 μm, zoom in panel scale bars, 1 μm. xz view along yellow dashed line. Scale bars, 2 μm. zoom in panel scale bars, 1 μm. Schematic shows the dimension of nanopattern. **c**, representative images showing adhesion morphology (top row: ꞵPS-GFP, bottom row: phalloidin) of isolated muscle cell adhesions on flat and nanotopographies with different width. **d**, Fluorescence intensity of ꞵPS-GFP. n = 32 cells for flat, n = 13 for 200 nm, n = 15 for 400 nm, n = 8 for 800 nm, n = 5 for 1600 nm, same for e. **e**, Fluorescence intensity of phalloidin. **f**, SptPALM of ꞵPS-mEos3.2 on nanotopography with 800 um width with DMSO control. Shaded area shows representative regions corresponding to bottom or top parts of the nanotopography (See illustration). **g**, SptPALM of ꞵPS-mEos3.2 on nanotopography with 800 um width with CK666 treatment. **h**-**i**, Diffusion fraction for bottom (h) and top (i). n = 11 cells for DMSO bottom, n = 9 for CK666 bottom, n = 12 for DMSO top, n = 11 for CK666 top, same for j-m. **j**-**k**, Diffusion Coefficient for bottom (j) and top (k). **l**-**m**, Step angle anisotropy for bottom (l) and top (m).

We then plated the muscles on engineered substrates with nanotopographies (600 nm height, 800 nm width) that mimic the geometry observed in mature MASs. This strongly promoted adhesion maturation and substantially increased the intensity of ꞵPS-GFP, as well as actin, inside adhesions (**Fig. 6b**, **Extended Data Fig. 17a, b**). The adhesions preferentially developed inside the nano-channels with higher intensity on the side walls (**Fig. 6b** and **Extended Data Fig. 17c, d**), but also formed clusters on the top and bottom surfaces (**Extended Data Fig. 17e**). Using nanotopographies with a fixed depth (600 nm) and varying widths, we observed a size-dependent promotion of adhesion formation on nano-structured substrates (200 nm, 400 nm, and 800 nm wide) compared to micro-structured (1600 nm wide) and flat substrates (**Fig. 6. c-e**). Thus, the ability to make protrusive structures confined within nanotopographies is essential for MAS maturation.

Since MAS maturation is correlated with the formation of ridge-like structures that could constitute diffusion traps, we then wondered whether nanotopography affects the diffusive behaviour of integrin. Tracking of ꞵPS-mEos3.2 showed increased immobilization, decreased diffusion coefficient and increased confinement on nanopatterns compared to on flat substrate (**Extended Data Fig. 17e-i**). The effects were more pronounced on the bottom surface compared to the top of the ridges (**Extended Data Fig. 17g-i**), suggesting that the confinement of integrin diffusion is primarily an effect of the protrusive nano-domains consisted of membrane enveloped adhesion complex and actin cytoskeleton. CK666 treatment reduced paxillin-GFP intensity on the nanopattern (**Extended Data Fig. 18a-b**), decreased ꞵPS-mEos3.2 immobile fraction, increased its diffusion coefficient, and decreased its confinement (**Fig. 6f-m** and **Extended Data Fig. 18c-n**), showing that the regulation of integrin diffusion by the nanotopography relies on Arp2/3-dependent actin polymerization, as in live tissue. Altogether these results suggest that the nanotopography of membrane and the actin cytoskeleton are essential for building long-lasting integrin adhesions in 3D tissue.

## Discussion

Here we demonstrate at the molecular level how membrane nanotopography induced by actin-based protrusions can directly influence the diffusion, immobilization, activation and mechanical connection of adhesion proteins and promote the formation of stable force resisting adhesions in developing tissue. The nanotopography induced diffusion trap provides a framework for local regulation of supramolecular organization that promote their aggregation through molecular crowding and increased probability for protein interactions ^64^. This complements and extends the molecular-clutch-based model of integrin adhesion ^45,65^ in a 3D tissue context where the motion of adhesion proteins and actin are affected by increased geometrical confinement. We envision the same principle can be applied to diverse 3D cellular and tissue structures in building stable force-resistant nano-domains that regulate protein diffusion, such as cadherin-mediated cell-cell adhesions and neuromuscular-junctions.

Although the importance of substrate and membrane nanotopography in promoting cell adhesion ^66^, migration ^67^ and differentiation ^68^ has been demonstrated, the molecular and biomechanical mechanisms are not clear. Membrane invaginations recruit actin regulators, including Arp2/3, BAR domain proteins, and integrins ^28,59,69^, and could mediate adhesion to protein fibres ^28^. Here we provide evidence for the regulation of integrin adhesion by Arp2/3-induced membrane protrusions. The constrains exerted by membrane curvature ^26^ and membrane proximal actin cytoskeleton ^70^ can limit the diffusion of membrane proteins and promote their clustering ^27^. The reduction of integrin diffusion and confinement inside diffusion nano-domains, observed in MASs, may help to concentrate adhesion proteins in a dynamic pool where a large portion of proteins remains diffusive but can be immobilized rapidly ^17,71^. This is consistent with the observation that a small fraction of talin molecules bear force *in vivo* ^72^, despite the fact that integrins and associated proteins are very concentrated in the adhesion. The confined movement of actin filaments and restricted diffusion of integrin adhesion proteins potentially create more possibilities for interaction and may lead to a stronger molecular connection *in vivo*.

In addition to regulating the molecular dynamics of integrins, the organization of the actin cytoskeleton and adhesive structure by nanotopographic patterns also provides an efficient configuration for force generation and mechanical stability. As the actin networks are sandwiched in between integrin adhesions, compressive force may help their engagement, thus limiting the optimal size of the nanodomains (**Fig. 6c-e**) to the submicron regime. In addition, the anchoring of actin protrusion on both sides, also promotes more efficient protrusions against apposing tissues, which in turn can exert force on the growing adhesions, thus forming a positive feedback loop. This configuration also results in force generated nearly parallel to the membrane which may help orient actin and adhesion proteins such as talin for mechanotransduction ^73–75^.

## Acknowledgements

We thank Christel Poujol and Magali Mondin from the Bordeaux Imaging Center (BIC) for technical assistance, Frank Schnorrer for helpful discussions, Sven Bogdan for gift of fly lines and Benoît Ladoux for helpful discussions and gift of reagents. We acknowledge financial support from the French Ministry of Research and CNRS, and the Human Frontiers Science programme (Grant RGP0009/2017). T.C. also received supported from the French government in the framework of the University of Bordeaux’s IdEx “Investments for the Future” program / GPR LIGHT. C.H.F.E. also received support from the University of Cambridge Career Support Fund. O.R., S.L.F. and G.G. are supported by the France BioImaging national infrastructure ANR-10-INBS-04. SLF acknowledges the funding from the European Union’s Horizon 2020 research and innovation programme under grant agreement No. 871124 Laserlab-Europe (JRA ALTIS), and from ANR (MSM project ANR 17-CE09-0040-02).

## Author contributions

T.C., C.H.F.E., N.B. and G.G. conceptualized the project and designed the experiments. T.C. performed embryo dissection, sptPALM and super-resolution imaging experiments, developed MATLAB and Python code and performed data analysis. C.H.F.E. and B.K. conducted fly genetics, generated transgenic fly lines. C.H.F.E. performed SIM imaging on fixed embryos. A.I., P.J. and S.L.F. developed and performed 3D ModLoc super-resolution imaging experiments. C-T.T., Y.Y. and B.C. designed and fabricated quartz nanopatterns. M.F. helped with fly husbandry and dissection. O.R. and B.K. performed pioneering experiments of sptPALM. T.C., N.B. and G.G. wrote the manuscript. All authors discussed the results and commented on the manuscript.

## Supplementary Information

### Materials and Methods

#### Fly genetics

The following Drosophila stocks were used in this study. A genomic rescue construct Paxillin-GFP ^1^ and Mys-GFP ^2^ was used to mark the muscle attachment sites. These flies were combined with UAS::Utrophin-GFP, UAS::Arp3-GFP, UAS::eWAVE-GFP and with new constructs made for this study, as described below: mys-mEos3.2, Fit1-mEos3.2, UASp::mEos3.2, UASp::mEos3.2-CAAX, and UASp::mEos3.2-Actin5C. The UAS constructs were expressed in the muscle by crossing to the driver *Mef2::Gal4*.

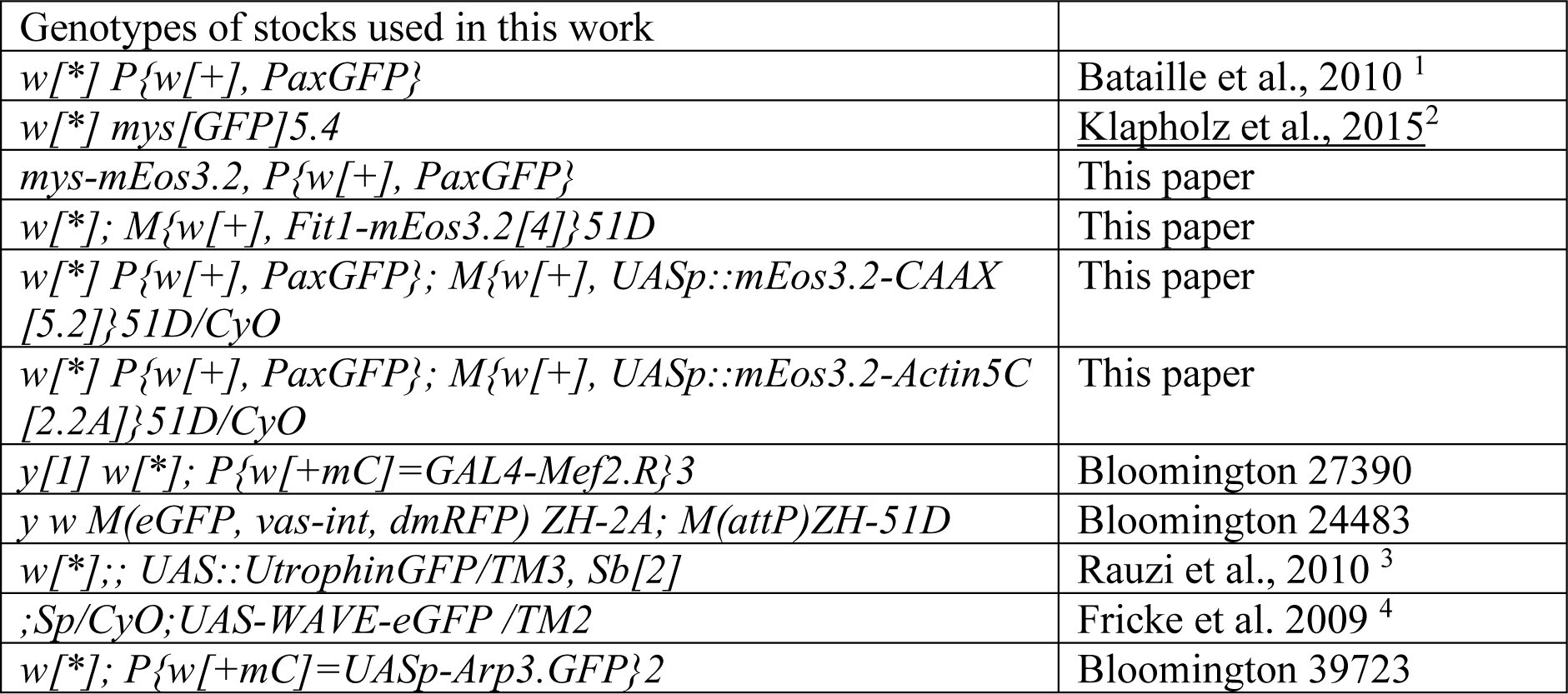

#### Transgenic constructs

##### Generation of tagged version of βPS (mys) and Fermitin1 (Fit1) with mEos3.2

An insertion of mEos3.2 into the endogenous locus encoding βPS, *(mys::mEos3.2)* was generated by homologous recombination, following the same procedure as for *mys-mCherry* ^5^. *Fit1-mEos3.2* is a genomic rescue construct of the *Fit1* gene, tagged with mEos3.2 internally within the F1 loop, with a linker of 4 serine residues at each junction. Thus, setting the A of the initiator ATG as 1, the construct extends from -1447 to 3994, with mEos3.2 inserted between bases 681 and 682 (amino acids 227 and 228). In a single copy this construct is able to restore viability and fertility to animals homozygous for a null allele of *Fit1.* The insertion of the fluorescent protein internally within Fit1 is essential, as constructs tagged at either the N- or C-terminus of Fit1 did not rescue the null *Fit1* mutant (unpublished observations).

##### Generation of UAS constructs

mEos3.2 was amplified from pmEos3.2-C1 plasmid with primers that added NotI and BamHI restriction sites to flank the coding sequence. The Kozak consensus sequence (CAAA), optimal for translation in *Drosophila*, was included just before the methionine initiation codon of mEos3.2. The CAAX sequence was added into the C-Terminus of mEos3.2 sequence by PCR amplification using a specific reverse primer that added the CAAX region just before the stop codon amino acid of mEos3.2.

The ubiquitous cytoplasmic actin, encoded by *Actin5C,* was tagged with mEos3.2 at the N-terminus as previously described ^6^. The coding sequence of mEos3.2 was amplifying by PCR and inserted into a UASp AttP vector, and the amplified Actin5C coding sequence appended to the mEos3.2 sequence with a GSGS linker sequence.

#### Generation of transgenic flies

Fit1mEos3.2, UAS::mEOs3.2, UAS::mEos3.2CAAX and UAS::mEos3.2Actin5C were inserted into embryos carrying an AttP docking site at cytological position 51D on the second chromosome, and on the X chromosome the φC31 integrase driven by the vasa promoter to express it within the germ line. Transformants were selected in the F1 generation by the *w+* eye colour, crosses were performed to remove the integrase and balance the stock. The new UAS lines were validated by combining with Mef2::Gal4 lines to drive expression in muscles and confirming photoconversion of mEos3.2.

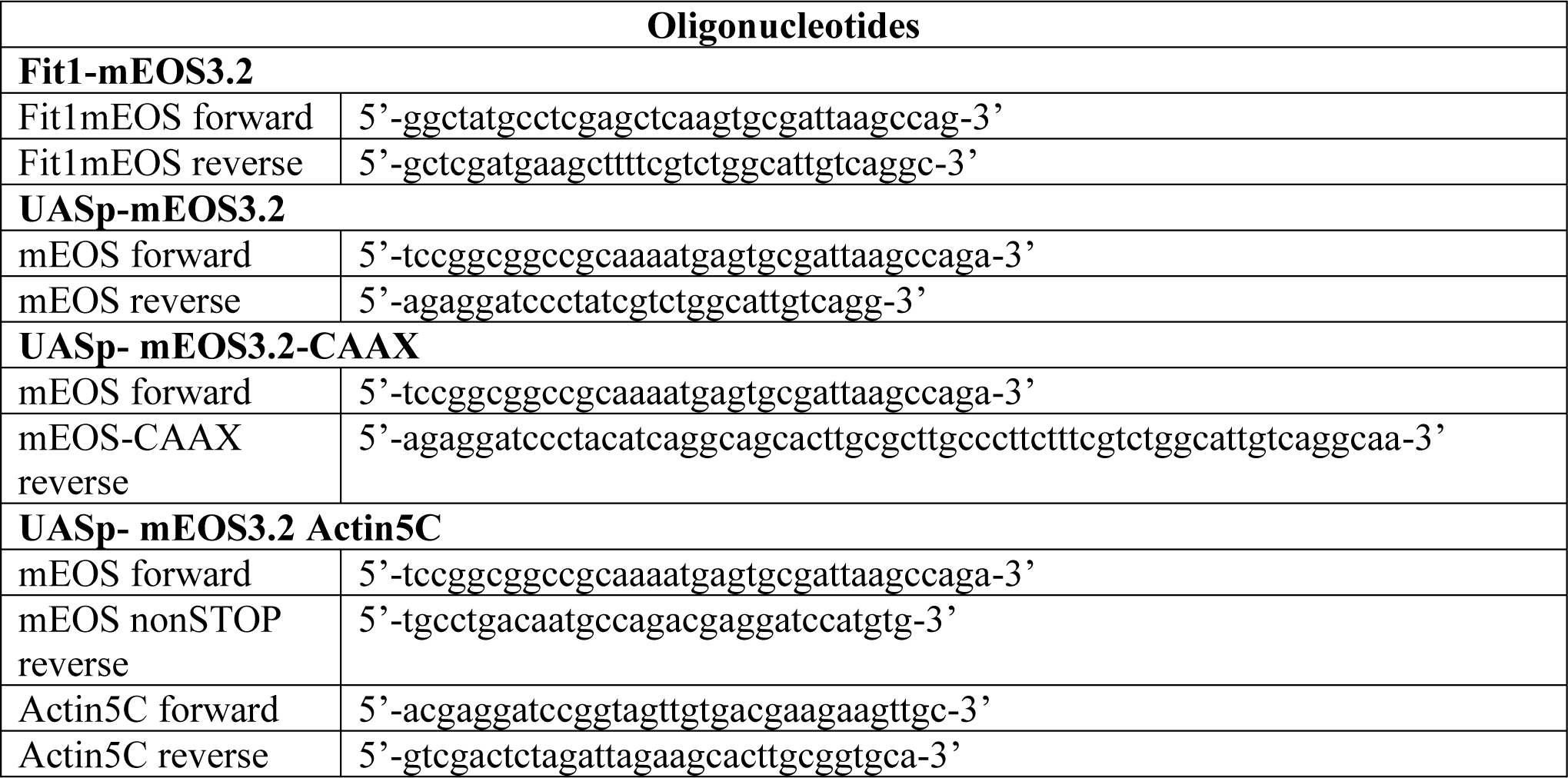

#### Embryo fixation and immunofluorescence microscopy

Embryos were collected on apple-juice agar plates spotted with fresh yeast. They were aged to stage 15-17 of development and processed for immunofluorescence using standard procedures. Embryos and first instar larvae were either imaged live or fixed. The sample were fixed in 4% formaldehyde and stained with phalloidin in PBST (PBS plus 0.5% bovine serum albumin and 0.3% Triton X-100). βPS-GFP was visualized directly in vivo or in fixed tissue. Comparison with the integrin signal was done using the anti-βPS monoclonal antibody (CG6G11, Developmental studies hybridoma bank). Actin visualization was performed using Rhodamine phalloidin and Alexa Fluor 647 phalloidin (Life Technologies) used at 1:500 and 1:50, respectively. Samples were scanned with a Leica SP8 confocal microscope using a 20X/0.75 NA objective for whole-embryo imaging or a 60X/1.40NA Oil objective for muscle attachments.

3D-SIM super resolution images were performed using a Zeiss Elyra 7 Lattice SIM, the system equipped with two sCMOS cameras allows the 2-color acquisition. The images were capture with 63x/1.4NA oil immersion lens. Raw data was reconstructed using ZEN software. Images were acquired with the optimal Z-step size covering several LT1-3 adhesion sites from the same embryo of a total of 22 wild type embryos. 3D visualization was done using Fiji software from a region of interest of 8-10 µm^2^ to cover a whole adhesion site.

#### Embryo fillet dissection

Flies were allowed to lay eggs on a grape juice agar plate at 25 °C for 2h and then aged at 18-22 °C. Early stage 16 embryos were selected based on gut morphology (to synchronize the timing of adhesion formation we started dissection at the point when the constrictions of the midgut segments begin to lose left-right symmetry). Embryos were briefly washed with 70% Ethanol and MQ water before dissection. The chorion was removed by rolling the embryo on a double-sided tape. During the dissection, a series of cuts were made with a tungsten needle along the mid-dorsal line from the posterior to the anterior that penetrate the vitelline membrane and epithelium while avoiding damaging the musculature. Embryos were then transferred and adhered to a 0.1% poly-L-lysine (Sigma) coated coverslip and flattened to form a fillet. The gut was removed to reduce autofluorescence and to prevent contractions that could interfere with imaging. Embryo fillets were washed 3 times with PBS and allowed to develop in Shields and Sang M3 insect medium (Sigma) supplemented with 2% FBS, 0.01 mg/ml Insulin (Sigma) and Pen/Strep.

#### Embryo fillet fixation and immunostaining

Embryo fillets were fixed in PEM buffer (80 mM PIPES, 5 mM EGTA, 2 mM MgCl_2_, pH 6.8) with 4% paraformaldehyde (PFA) and 0.2% glutaraldehyde for 10 min at 37 °C. Subsequently, the embryo fillets were washed 3 times with PBS and permeabilized in PBST (0.2% Triton X-100) for 5-7 days at 4 °C, before blocking in PBST with 3% BSA overnight at 4 °C. The following primary and secondary antibodies was used: anti-βPS monoclonal antibody (CG6G11, Developmental studies hybridoma bank) (1:15), CF680 goat anti-mouse IgG (Biotium) (1:200). F-actin was visualized with phalloidin-AF647 (Life Technologies) (1:100).

#### Confocal and SIM microscopy of live embryo fillet

Confocal and SIM acquisitions were performed on a spinning disk confocal microscope (Leica DMI8, 100X 1.49 N.A. oil immersion objective, sCMOS Camera Photometrics Prime 95B). For MAS development over several hours, one z-stack of 10 μm with 0.2 μm step size was acquired at 10 minutes per frame for 6-8 hours. For fast dynamics of integrin clusters and actin puncta, one z-stack of 5 μm with 0.2 μm step size was acquired at 10 seconds per frame for 5-10 minutes. Super-resolution SIM images were acquired with the Live-SR module.

#### Spectral demixing STORM

Spectral demixing STORM ^7^ acquisition was performed on an Abbelight SAFe 360 module installed on and inverted microscope (Nikon Ti) with a CFI Apo TIRF 100X oil 1.49 NA objective using HiLo illumination. Samples stained with AF647 and CF680 was imaged with a 640 nm laser. Fluorescence was collected with a dichroic (Dio02-R561-25×36, Semrock) and an emission filter (FF01-432/515/595/730-25, Semrock). The fluorescence was subsequently separated by a dichroic (FF699-Fdi01-t3-25×36, Semrock) and collected by two cameras (Hamamatsu ORCA-Flash 4.0). The images were analyzed with the Abbelight Neo software as per the manufacturer’s instructions. Demixing of the localizations based on the ratio (0-1) of photons from the two cameras were calculated and the ratios 0-0.4 were assigned to the AF647 channel while the ratios 0.6-1 were assigned to the CF680 channel. Ratios between 0.4 and 0.6 were rejected.

#### 3D ModLoc STORM

For 3D ModLoc STORM ^8^, the sample was excited by a 639nm laser (Genesis MX 639, 1W, Coherent) and imaged through a Nikon APO TIRF 60x 1.49 NA oil immersion objective lens mounted on an inverted microscope (Eclipse Ti, Nikon). The fluorescence was collected with a dichroic (Di03-R635-t1-25×36, Semrock) and an emission filter (BLP01-635R-25, Semrock) on four quadrants of a 512×512 pixel EMCCD camera (iXon3, Andor). An EOM (EO-PM-NR-C1, Thorlabs) was used for the phase modulation at the excitation whereas a Pockels cell (CF1043-20SG-500/700, Fast pulse) performed the demodulation at the detection. 60 cycles of modulation/demodulation were performed during the acquisition time of one image (50 ms). Typically between 20000 and 30000 images were acquired to reconstruct the 3D super-resolved images. Drosphila embryo fillet samples were imaged 5-10 μm in depth with a z range of 1 μm.

#### sptPALM

Embryo fillets or isolated muscle cells were imaged at 26 °C in a Ludin chamber (Life Imaging Services) on and inverted microscope (Nikon Ti) with a CFI Apo TIRF 100X oil 1.49 NA objective with HiLo illumination. A 561nm laser (Cobolt, 20 mW out of objective) was used to excite mEos3.2 while simultaneously photo-activated with a 405nm laser (Omicron, 5mW out of objective). The laser power of the 405nm laser was adjusted for each sample to control the density of activated mEos3.2 and to ensure a spatial temporal separation of single molecule signals enabling localization and tracking. Fluorescence of mEos3.2 was collected with a Di01-R561 dichroic and FF01-617/73 emission filter (Semrock) and an EMCCD camera (Evolve, Photometrics) Image streams were acquired with Metamorph software (Molecular Devices) at 50Hz (20ms per frame) and approx. 10,000 frames were acquired for each set of experiment. A reference image of Pax-GFP is acquired before and after the stream with a GFP filter cube (excitation filter: FF01-472/30, dichroic: FF-495Di02, emission filter: FF02-520/28; Semrock).

#### Single-molecule tracking analysis

Single molecule fluorescence from sptPALM images were localized with the PALMtracer software ^9^ where a wavelet based method is used to identify the spots followed by gaussian fitting of the point spread function to localize the 2D position of the fluorophore. Single molecule localization from consecutive frames are then linked with a maximum distance of 3 pixels per frame to track diffusing molecules. Trajectories lasting at least 10 frame were processed with a custom written MATLAB code for mean squared displacement (MSD) analysis. The MSD is calculated as:

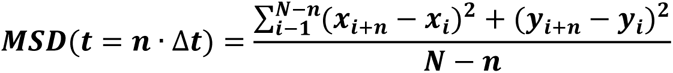

Where x_i_ and y_i_ are the 2D coordinates of the molecule at time ***i*** · Δ***t***. The proteins in the cell can exhibit anomalous diffusion at longer time scale but at shorter time scale will appear diffusive with a close to linear MSD curve. We fit the first 4 points of the MSD and take the slope as the apparent diffusion coefficient D. In order to differentiate the diffusive and immobile trajectories, we use two parameters D_thresh_ and R_conf_. D_thresh_ = (2.355 σ) ^2^/(4×4×0.02s) is the minimum D observable limited by the localization uncertainty σ of the imaging system. R_conf_ is the confinement radius by fitting the first 80% of the MSD curve with the following:

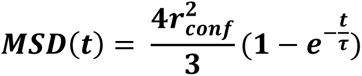

where 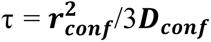. We characterize a trajectory as immobile if its D< D_thresh_ and R_conf_ <3σ. For each set of experimental conditions, the localization uncertainty σ is characterized with the ThunderSTORM plugin in Fiji, and the resolution is determined as 2.355 σ (88.3 nm for embryo fillet and 93.8 nm for Curi Bio nano-pattern). The respective D_thresh_ is 0.0243 µm^2^/sec for embryo fillet and 0.0275 µm^2^/sec for Curi Bio nano-pattern, and is represented by the gray area in the diffusion coefficient distribution graphs.

In order to quantify the confinement of the trajectories, we measure the step angle between consecutive steps of only diffusive trajectories and only for steps larger than the resolution limit. From the distribution of the angles we then calculate the step angle anisotropy = f_150-210_/60×360, where f_150-210_ is the fraction of angles between 150 and 210 degrees.

#### Drug treatment and Mn^2+^ stimulation

Chemical inhibitors were administered at the following concentration: Latrunculin A (Enzo Life Sciences), 10 mM; CK666 (Tocris), 250 μM; Swinholide A (Abcam), 100 nM; Y-27632 (Sigma), 100 μM. Due to the photo-sensitivity of Y-27632, the treatment was performed in a dark environment prior to imaging.

A brief stimulation with 5 mM Mn^2+^ was performed in Ringer’s solution (150 mM NaCl, 5 mM KCl, 2 mM CaCl_2_, 2 mM MgCl_2_, 10 mM Hepes and 2 g/L Glucose, pH 7.4) at different developmental stages for 15 min before sptPALM imaging. This was followed by exchanging to culture medium after imaging to allow embryo fillet development.

#### F-actin movement measurement

Female flies carrying UAS-Actin-mEos3.2 and Paxillin-GFP were crossed with males carrying Mef2-GAL4. Embryos were collected and dissected for fillets. Expression of Actin-mEos3.2 was confirmed with photo-activation of 405 laser before imaging. Single molecule dynamics of Actin-mEos3.2 was imaged with 2 Hz frame rate and 500 ms exposure time with a 561 laser and HiLo illumination. This imaging frequency excludes the fast diffusing actin monomers and tracks only actin within slow moving filaments. Localization and trajectories linking were performed with PALMtracer. Trajectories were classified into mobile and stationary with MSD analysis using the following parameters: D_thresh_ = 0.000975 µm^2^/sec, σ = 0.0375 µm. A custom written MATLAB routine was used to calculate the actin movement speed and distribution of angles. The movement speed was defined as the distance travelled from the beginning to the end of the track divided by duration of track. Movement angle was based on the vector from the beginning of the track to the end of the track. To characterize the type of movement of actin trajectories, the mobile trajectories were fitted with the power law 4Dt^α^. Tracks with α<0.8 were classified as sub diffusive, 0.8<α<1.2 were classified as Brownian diffusion and α>1.2 were classified as directed motion ^10^.

#### Actin puncta and integrin foci tracking

Adhesions in embryo fillets expressing Utrophin-GFP or ꞵPS-GFP were imaged at different developmental stages at 10 sec/frame with a spinning disk confocal microscope (Leica DMI8, 100X 1.49 N.A. oil immersion objective, sCMOS Camera Photometrics Prime 95B). Actin and integrin images were first background subtracted with a 3 pixel radius rolling ball to reveal dynamic puncta from stable structures. Puncta were detected and tracked with Fiji TrackMate plugin. Speed of the puncta was calculated as instantaneous speed from consecutive frames.

#### Soft PDMS gel fabrication and functionalization

Soft PDMS gel base (Dowsil CY-52-276-A) and curing agent (Dowsil CY-52-276-B) were mixed at ratios 1.2:1, 1:1 and 1:1.2 to produce gels with different rigidities of approx. 3 kPa, 15kPa and 35 kPa respectively ^11^. One drop of mixed gel was deposited on an 18mm coverslip and spin coated at 2000 rpm for 30 sec to produce a thin layer of approx. 30-50 µm. The gels were cured at 80 °C for 2h. To functionalize the surface of the gel, 5% (3-aminopropyl) triethoxysilane (APTES) (Sigma) was deposited on the gel in ethanol for 5 min, washed 1X with ethanol and subsequently cure at 70 °C for 30 min. Then 10 μg/ml Human Fibronectin (Sigma) was coated for 1h at 37 °C before washing and seeding of cells.

#### Fabrication of quartz nanopattern

Nanopatterned gratings were fabricated on cleared fused quartz coverslips measuring 18mm in diameter and 0.2 ± 0.05mm in thickness, obtained from Technical Glass Product Inc. Prior to usage, the coverslips underwent a thorough cleaning process involving acetone and isopropanol with sonication to remove any particles. Subsequently, the cleaned coverslips were blow-dried and heated at 180 degrees on a hotplate to ensure complete dryness. We then spin-coated a 275 nm layer of 9% CSAR 62 Ebeam resist onto the substrate at 2000 rpm and baked it at 180 degrees for 3 minutes. On top of this resist layer, we spin-coated a 100 nm conductive layer of Electra 92 at 1000 rpm and baked it for an additional 3 minutes. Using the Raith Voyager lithography system, we patterned the gratings with various dimensions (200 nm, 400 nm, 800 nm, 1600 nm) and pitches, incorporating multiple biases to compensate for proximity effects. Following lithography exposure, the patterned substrate was immersed in DI water to remove the Electra 92 conductive layer and then developed in xylene for 45 seconds to reveal the nanopatterns. We evaporated a 50 nm Cr layer onto the substrate as the masked and performed plasma etching using the PlasmaTherm Versaline LL ICP Dielectric Etcher for 600 nm. The Cr layer was removed with a Cr etchant for 30 minutes, leaving us with the desired nanopatterned gratings.

#### Isolation of embryonic muscle cells and *in vitro* culture

Mid stage 16 Drosophila embryos (after symmetry breaking of the gut, roughly corresponding to 3h *ex vivo*) were harvested and dissected as described before to produce a flat embryo fillet attached to a poly-L-lysine (Sigma) coated coverslip. Excess tissue including the gut, CNS and trachea were removed with a dissection needle. Other isolated cells were removed by gentle pipetting. The remaining musculature and epithelium attached to the cuticle were then washed 3X with PBS. A short treatment of 37 °C pre-warmed trypsin (without EGTA, Invitrogen) was administered for 3 min at room temperature to loosen the muscle tissue before exchanging to insect medium with 10% FBS. The muscle cells were then carefully teased out with a dissection needle and transferred with a 20 μl pipette tip to petri dishes with different substrates. The muscle cells were allowed to attach and develop adhesions for 1.5 h before imaging.

For experiments in Fig. 6c-e, a quartz coverslip with fabricated nanopatterns of different sizes were used as substrate. The surface was treated with oxygen plasma for 5 min before coating with 10 ug/ml Human Fibronectin (Sigma) for 1h at 37 °C and overnight at 4 °C. After cell seeding and imaging, the coverslip was washed with 10% SDS and subsequently immersed in concentrated sulfuric acid for 3 days to remove organic matter before washing with MQ water and reuse. For experiments in Fig. 6a-b, f-m and Extended Data Fig. 17 & 18, a commercial polymer nano-pattern or flat surface (Curi Bio) was used.

#### Image processing

Fluorescence images displayed in the figures were background subtracted and adjusted for better contrast. Fluorescence intensity measurements were done using raw images, where camera offset measured from closed shutter images were first subtracted.

ModLoc super-resolution localization data was first processed with a custom Python code to perform DBSCAN (scikit-learn) clustering analysis. Then, localizations outside clusters (represented by grey dots in Fig. 1d 2D density) were filtered out and 3D surface models were generated with ChimeraX software, a random colour is assigned to each cluster.

3D fluorescence images were rendered with Napari, and movies were generated with the animation plugin with key frames.

#### Statistical analysis

Data presented in bar and line graphs represent mean ± s.e.m. unless otherwise specified. Data presented in boxplots represent median±25/75 percentiles with range and statistical outliers marked. Data exclusion is based on sample size (minimum number of trajectories >100 per region of interest and length of individual trajectories ≥10) while statistical outliers were not excluded from analysis. Statistical tests were conducted with MATLAB. Two-tailed student’s t-tests were performed to compare data sets with n.s. representing p≥0.05, * representing p<0.05, ** representing p<0.01 and *** representing p<0.001. Normality of the data was tested by Anderson-Darling test with a confidence interval of 0.05.

### Extended Data Figures

**Extended Data Figure 1.**
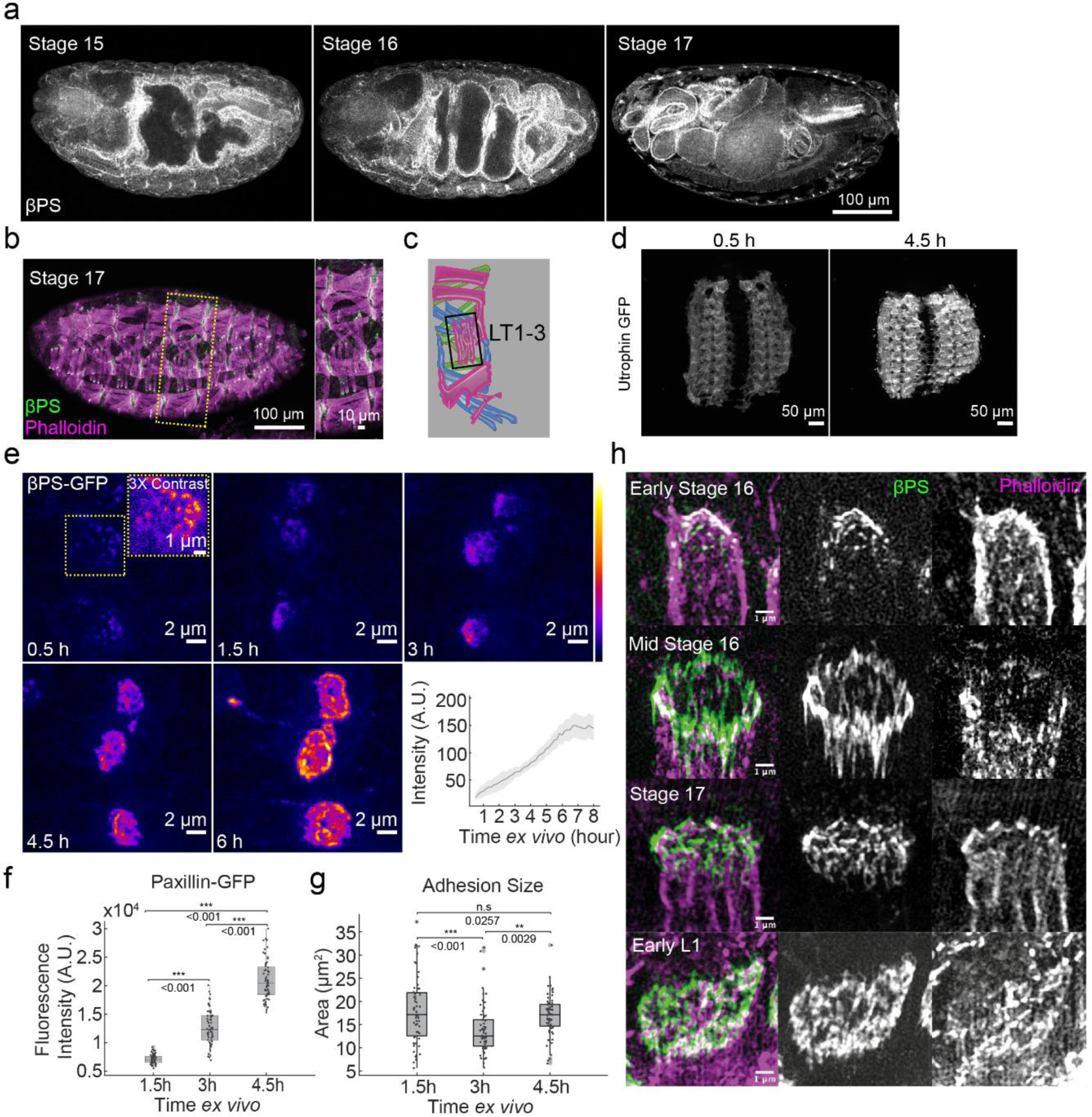
MAS adhesion development in Drosophila embryo and embryo fillet. **a**, Drosophila embryos from stage 15, 16 and 17 stained for βPS Integrin where typical gut morphology can be observed. Scale bar, 100 μm. **b-c**, Lateral view and schematic of a stage 17 embryo stained for βPS Integrin (green) and Actin (Phalloidin in green) showing the position and adhesion shape of the lateral-transverse muscles (LT1-3). Scale bar, 100 μm. Inset scale bar, 10 μm. **d,** Utrophin-GFP images showing development of muscle actin in embryo fillet. Scale bars, 50 μm**. e,** Spinning disk confocal images showing the morphology of LT1-3 muscle attachment site adhesions during development in embryo fillets. Scalebars, 2 μm. Integrin (βPS-GFP) enrichment can be observed in the adhesion site during development and quantification of fluorescence intensity (mean ± s.d.), n = 11 embryos. **f**, Quantification of the enrichment of Paxillin-GFP in developing MAS. n = 72 embryos for 1.5h, n = 77 for 3h, n = 76 for 4.5h. **g**, Quantification of adhesion size in developing MAS. n= 72 embryos for 1.5h, n = 77 for 3h, n = 76 for 4.5h. **h**, Morphology of LT muscle adhesions *in vivo* from early stage 16 embryos to early L1 larvae. Scale bars, 2 μm

**Extended Data Figure 2.**
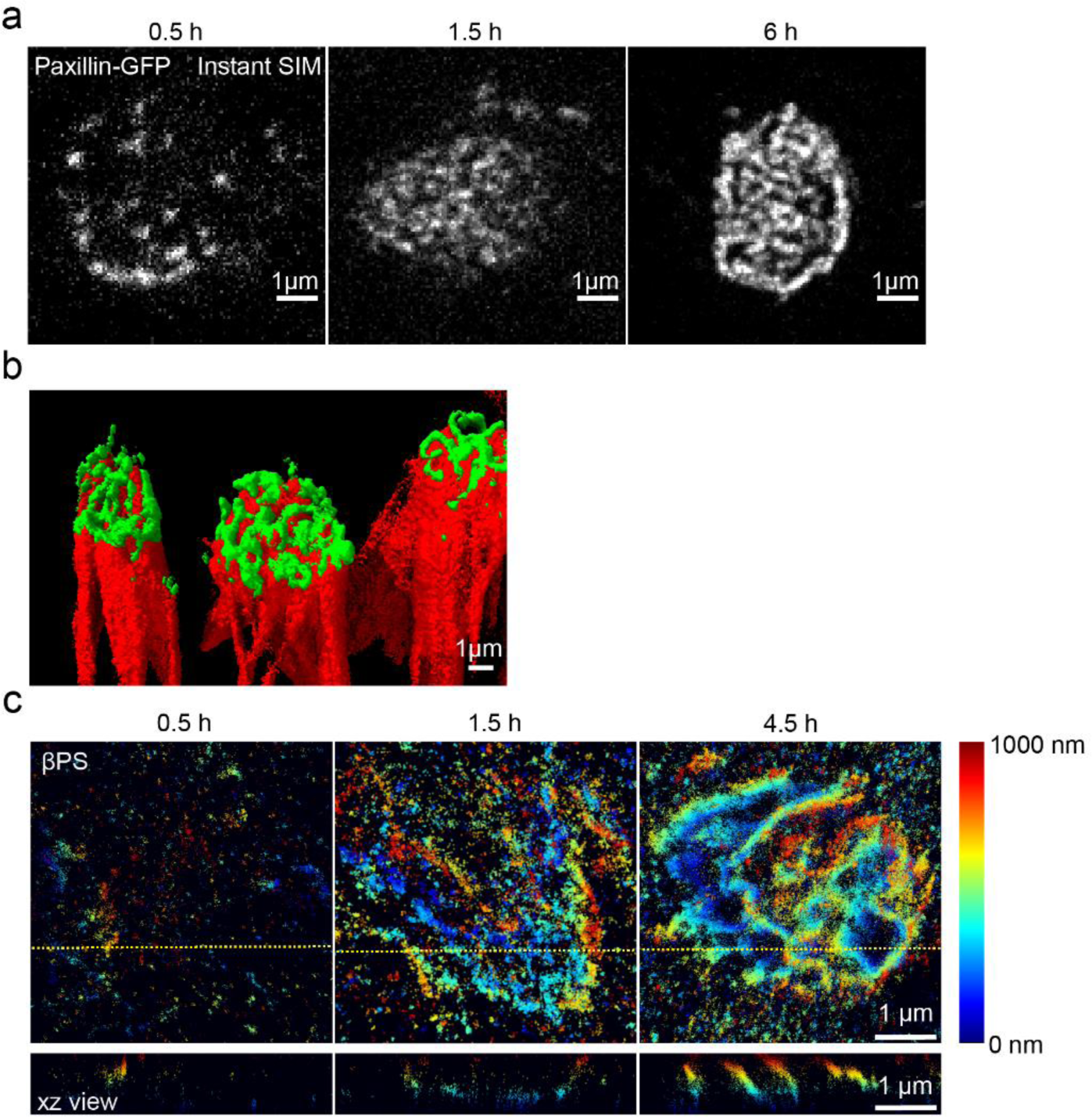
Super-resolution microscopy reveals adhesion structure in developing muscle attachment. **a**, SIM imaging of developing adhesions. Scale bars, 1 μm. **b**, 3D reconstruction of adhesion, Phalloidin: red, βPS-integrin: green. Scale bar, 1 μm. **c**, Representative 3D Modloc super-resolution images showing distribution of βPS-integrin in different developmental stages. Xz view shows cross section along the yellow dashed line. Scale bars, 1 μm.

**Extended Data Figure 3.**
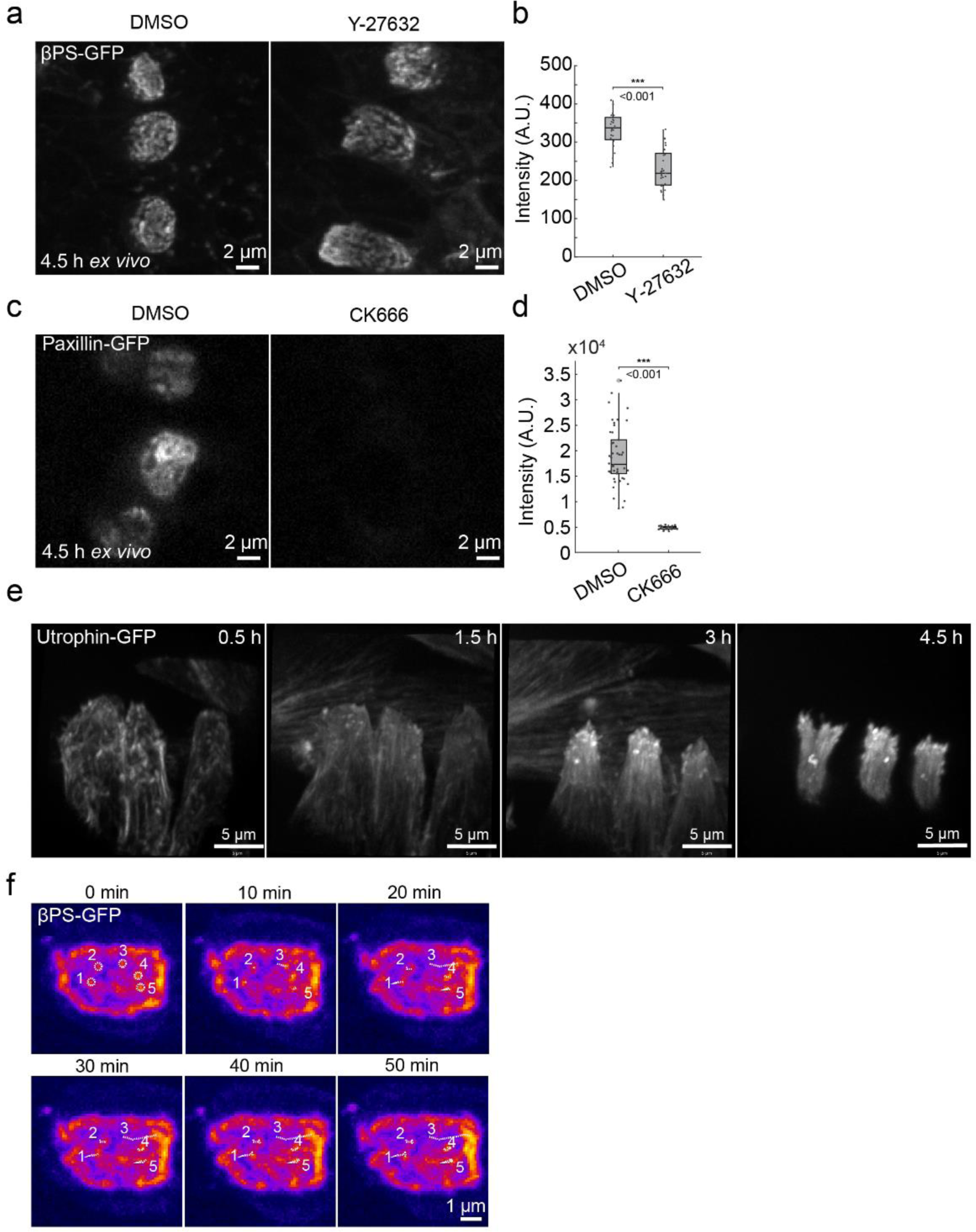
Arp2/3-dependent actin dynamics drive adhesion development. **a**, βPS-GFP images of 4.5h adhesions in DMSO control and Y-27632 treated embryo fillets. Scale bars, 2 μm. **b**, Quantification of βPS-GFP fluorescence intensity. n = 24 sets of adhesions for DMSO, n = 33 for Y-27632. **c**, Paxillin-GFP images of 4.5h adhesions in DMSO control and CK666 treated embryo fillets. Scale bars, 2 μm. **d**, Quantification of paxillin-GFP fluorescence intensity. n = 46 sets of adhesions for DMSO, n = 21 for CK666. **e**, 3D reconstruction of Utrophin-GFP in developing adhesions. Scale bars, 5 μm. **f,** Time-lapse images in a developing adhesion showing integrin cluster dynamics in a developing adhesion. White circles, initial positions of integrin cluster. White dashed lines, traces of moving integrin clusters. Scale bars, 1 μm.

**Extended Data Figure 4.**
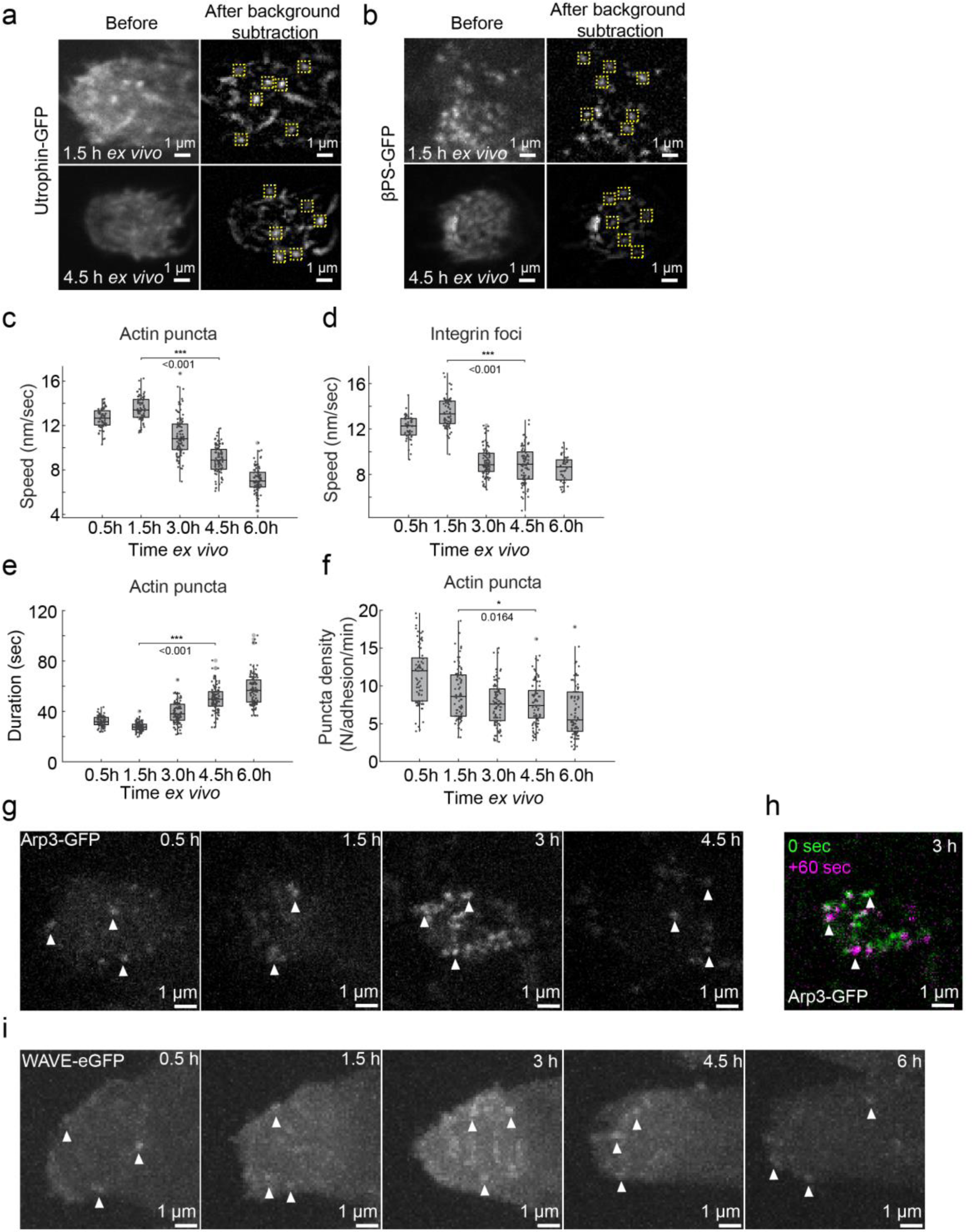
Dynamics of actin puncta and integrin clusters during adhesion development. **a**, Background subtraction of Utrophin-GFP images and detection of actin puncta (yellow boxes). Scale bars, 1 μm. **b**, Background subtraction of βPS-GFP images and detection of integrin foci (yellow boxes). Scale bars, 1 μm. **c**, Movement speed of actin puncta. n = 69 adhesions for 0.5h, n = 81 for 1.5h, n = 89 for 3h, n = 85 for 4.5h, n = 84 for 6h. **d**, Movement speed of integrin foci. n = 52 adhesions for 0.5h, n = 73 for 1.5h, n = 87 for 3h, n = 90 for 4.5h, n = 42 for 6h. **e**, Duration of actin puncta. n = 87 adhesions for 0.5h, n = 101 for 1.5h, n = 110 for 3h, n = 107 for 4.5h, n = 114 for 6h. **f**, Density of actin puncta. **g**, Representative images showing Arp3-GFP puncta inside developing adhesions. Scale bars, 1 μm. **h**, Arp3-GFP puncta dynamics in developing adhesion (3h *ex vivo*). Scale bars, 1 μm. **i**, Representative images showing WAVE-eGFP puncta inside developing adhesions. Scale bars, 1 μm.

**Extended Data Figure 5.**
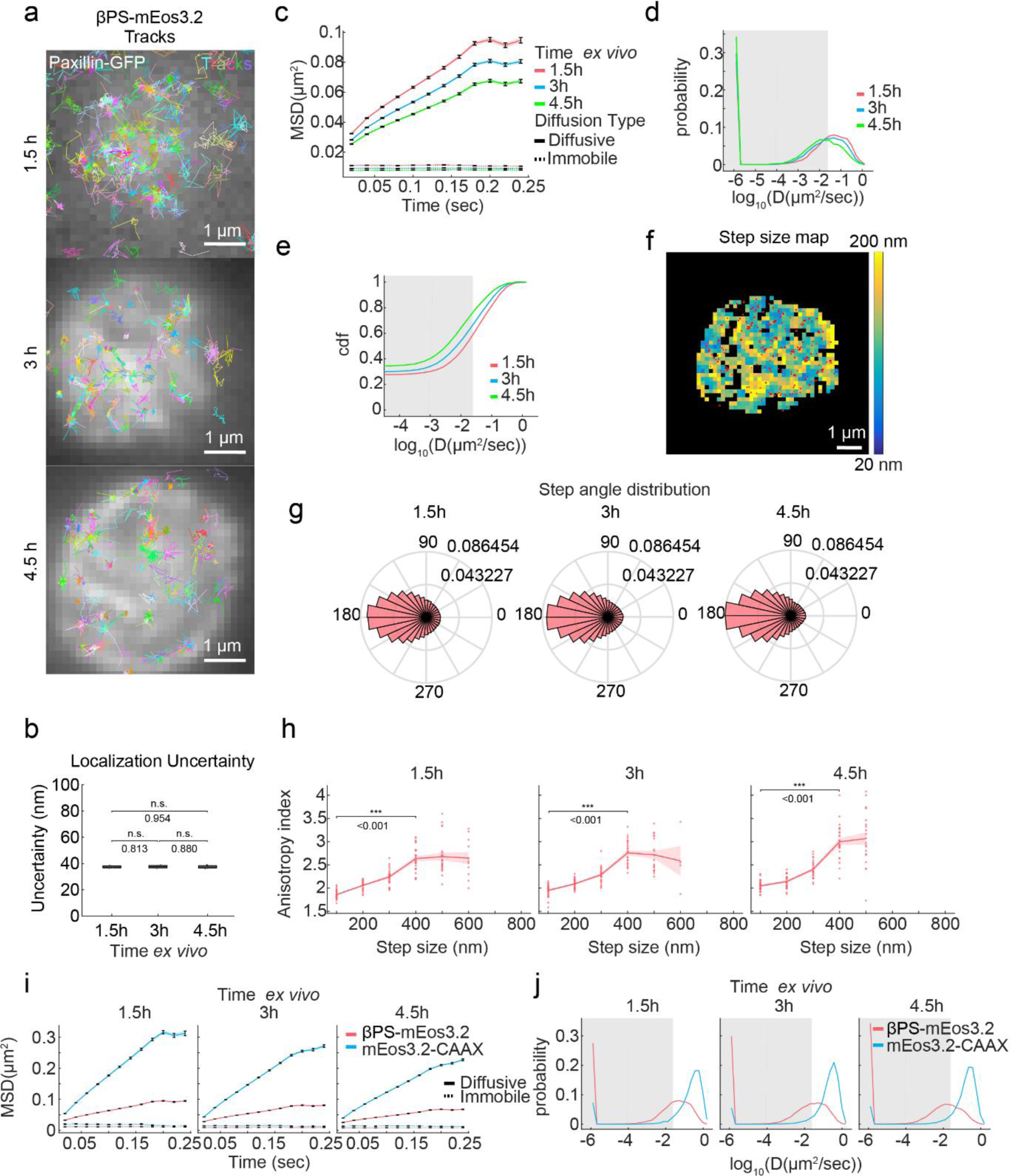
sptPALM tracking of βPS-mEos3.2 in developing adhesions. **a**, sptPALM tracking of βPS-mEos3.2, tracks are plotted with a random color. Scale bars, 1 μm. **b**, Localization uncertainty measured in live samples of βPS-mEos3.2 in embryo fillets at 1.3h, 3h and 4.5h *ex vivo*. n = 4 independent experiments each containing 5-10 embryos with thousands of trajectories for 1.5, 3h and 4.5h. **c**, Mean square displacement (MSD) of diffusive and immobile βPS-mEos3.2 tracks in 1.5h, 3h and 4.5h adhesions. **d**, Diffusion coefficient distribution of βPS-mEos3.2 tracks (probability). **e**, Diffusion coefficient distribution of βPS-mEos3.2 tracks (cdf). **f**, Step size map in 4.5h adhesion, red dots represent hot spots of immobilization. Scale bar, 1 μm. **g**, Step angle distribution. **h**, dependence of step angle anisotropy on step size. **i**, Comparison of βPS-mEos3.2 and mEos3.2-CAAX MSD. **j**, Comparison of βPS-mEos3.2 and mEos3.2-CAAX Diffusion coefficient distribution (probability).

**Extended Data Figure 6.**
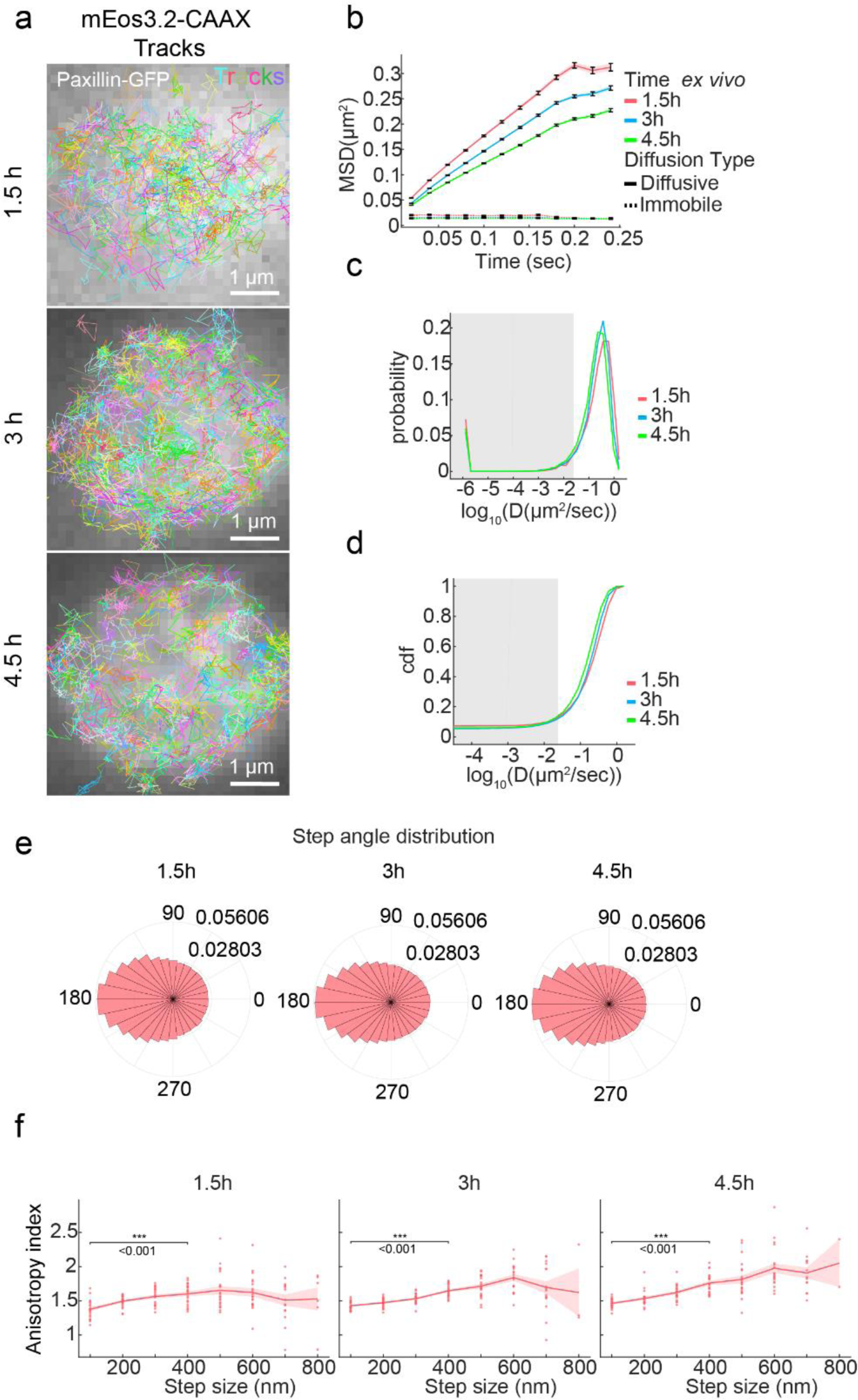
sptPALM tracking of mEos3.2-CAAX in developing adhesions. **a**, sptPALM tracking of mEos3.2-CAAX. Left column: tracks are characterized into diffusive (green) or immobile (red); Right column: tracks are plotted with a random color. Scale bars, 1 μm. **b**, Mean square displacement (MSD) of diffusive and immobile mEos3.2-CAAX tracks in 1.5h, 3h and 4.5h adhesions. **c**, Diffusion coefficient distribution of mEos3.2-CAAX tracks (probability). **d**, Diffusion coefficient distribution of mEos3.2-CAAX tracks (cdf). **e**, Step angle distribution of mEos3.2-CAAX tracks. **f**, dependence of step angle anisotropy on step size.

**Extended Data Figure 7.**
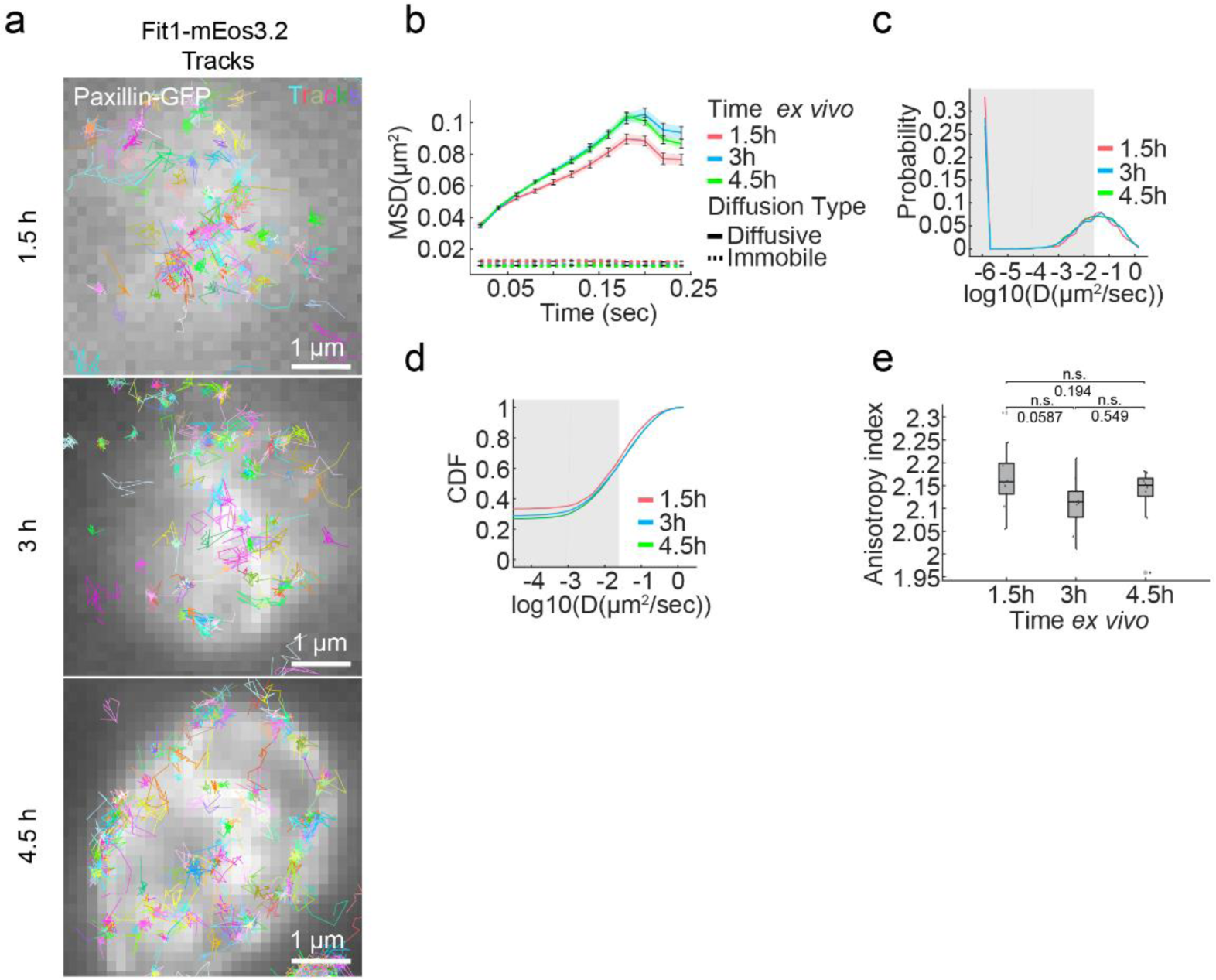
sptPALM tracking of Fit1-mEos3.2 in developing adhesion site. **a**, sptPALM tracking Fit1-mEos3.2. Left column: reconstructed PALM images; centre column: tracks are characterized into diffusive (green) or immobile (red); Right column: tracks are plotted with a random color. Scale bars, 1 μm. **b**, Mean square displacement (MSD) of diffusive and immobile fermitin1-mEos3.2. **c**, Diffusion coefficient distribution of fermitin1-mEos3.2 tracks (probability). **d**, Diffusion coefficient distribution of fermitin1-mEos3.2 tracks (cdf). **e**, Step angle anisotropy. n= 10 embryos for 1.5h, 3h and 4.5h.

**Extended Data Figure 8.**
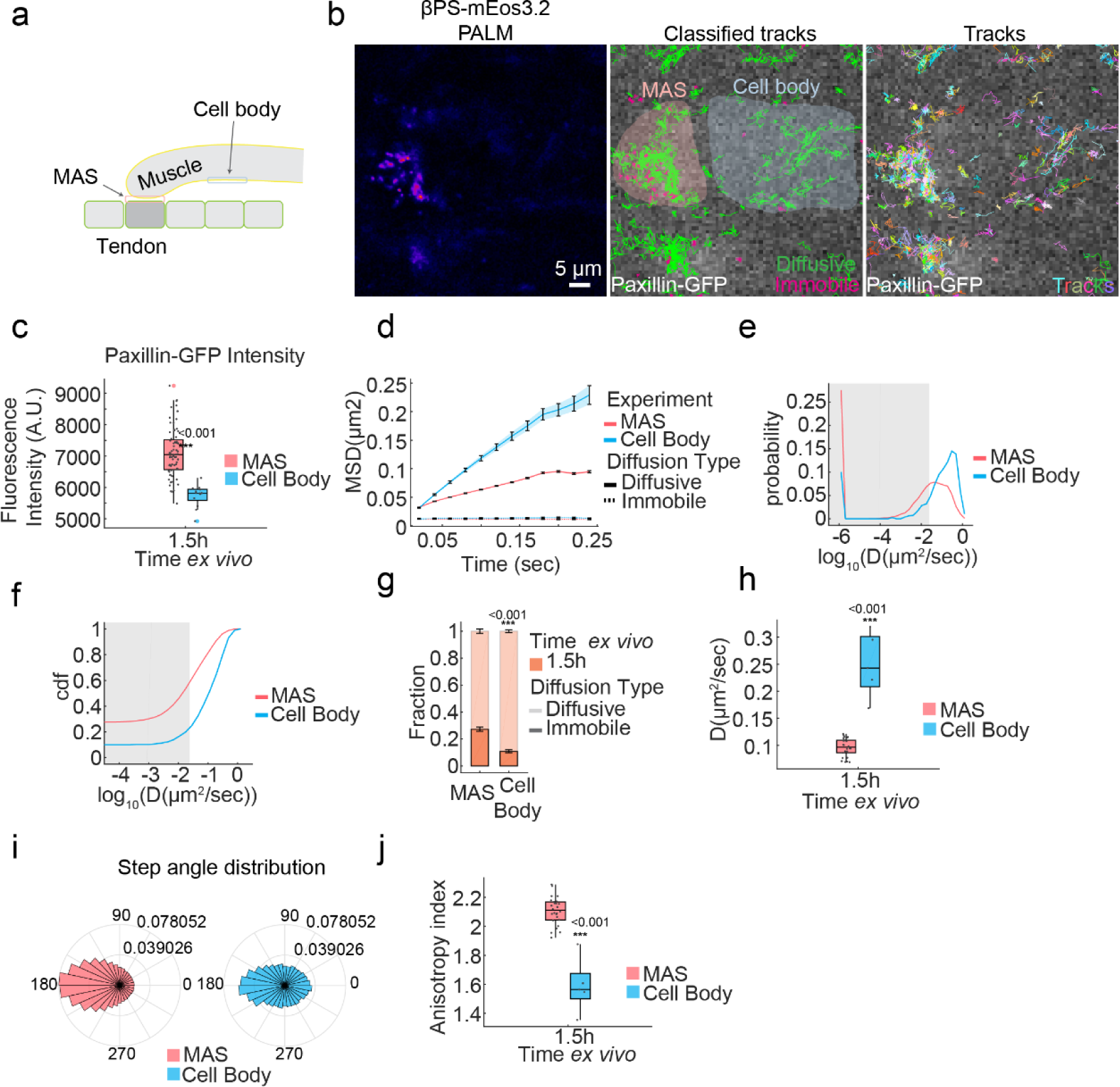
sptPALM tracking of βPS-mEos3.2 in the cell body. **a**, Schematic showing location of imaging for cell body and MAS. **b**, sptPALM tracking βPS -mEos3.2. Left column: reconstructed PALM images; centre column: tracks are characterized into diffusive (green) or immobile (red); Right column: tracks are plotted with a random colour. Scale bar, 5 μm. **c**, Paxillin-GFP fluorescence intensity quantification. n=72 adhesions for MAS, n=13 for cell body. **d**, Mean square displacement (MSD) of diffusive and immobile βPS-mEos3.2. **e**, Diffusion coefficient distribution of βPS-mEos3.2 tracks (probability). **f**, Diffusion coefficient distribution of βPS-mEos3.2 tracks (cdf). **g**, Fraction of Immobile and diffusive tracks. n= 26 embryos for MAS, n=5 for cell body. **h**, Diffusion coefficient. n= 26 embryos for MAS, n=5 for cell body. **i**, Step angle distribution. **j**, Step angle anisotropy. n= 26 embryos for MAS, n=5 for cell body.

**Extended Data Figure 9.**
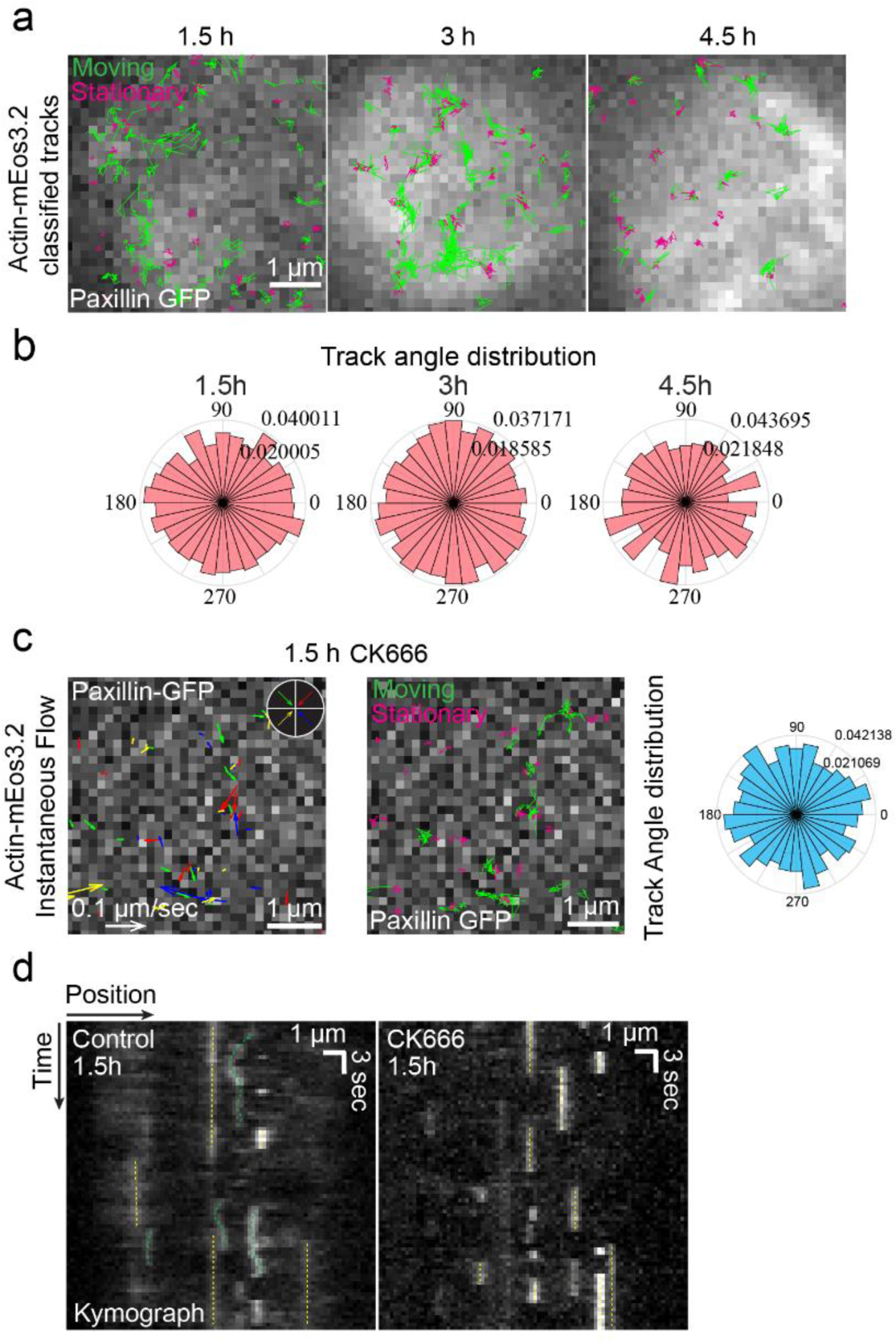
F-actin motion in developing adhesion. **a**, sptPALM tracking of actin-mEos3.2. Scale bar, 1 μm. **b**, Angle distribution of moving actin-mEos3.2 tracks (begging to end of track). **c**, F-actin motion in adhesion treated with CK666. Left panel scale bars, left: 0.1 μm/sec, right: 1 μm. Centre panel scale bar, 1 μm. Right panel, angle distribution of moving tracks. **d**, Kymograph showing stationary (yellow dashed line) and moving (green dashed line) actin-mEos3.2 dots in control and CK666 treated adhesions. Horizontal scale bar, 1 μm. Vertical scale bar, 3 sec.

**Extended Data Figure 10.**
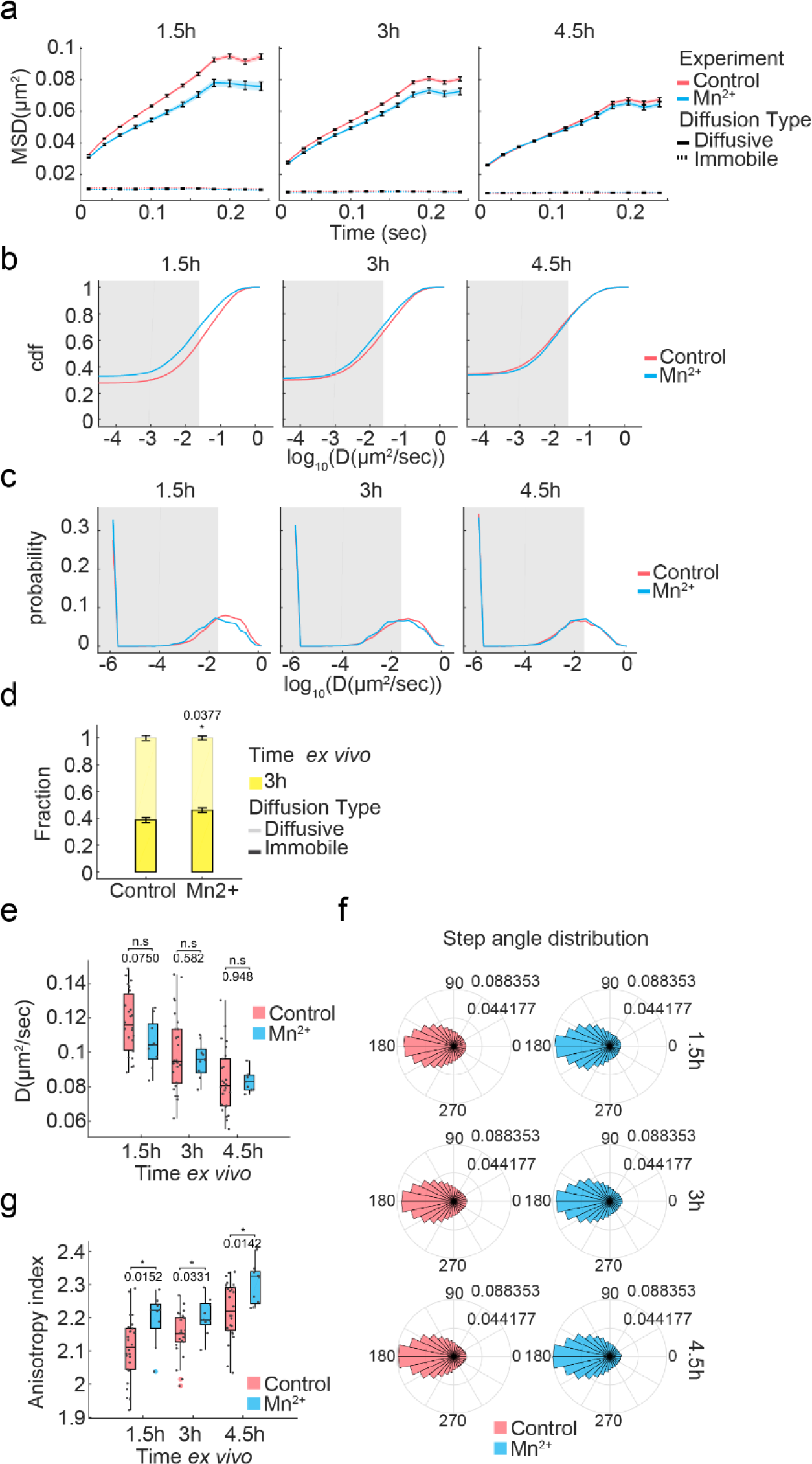
sptPALM tracking of βPS-mEos3.2 in developing adhesion with Mn^2+^ stimulation. **a**, Mean square displacement (MSD) of diffusive and immobile βPS-mEos3.2. **b**, Diffusion coefficient distribution of βPS-mEos3.2 tracks (probability). **c**, Diffusion coefficient distribution of βPS-mEos3.2 tracks (cdf). **d**, Fraction of Immobile and diffusive tracks. n= 26 embryos for control, n=9 for Mn^2+^. **e**, Diffusion coefficient. n= 26 embryos for control 1.5h, 3h and 4.5h, n=9 for Mn^2+^ 1.5h, 3h and 4.5h. **f**, Step angle distribution. **g**, Step angle anisotropy. n= 26 embryos for control 1.5h, 3h and 4.5h, n=9 for Mn^2+^ 1.5h, 3h and 4.5h.

**Extended Data Figure 11.**
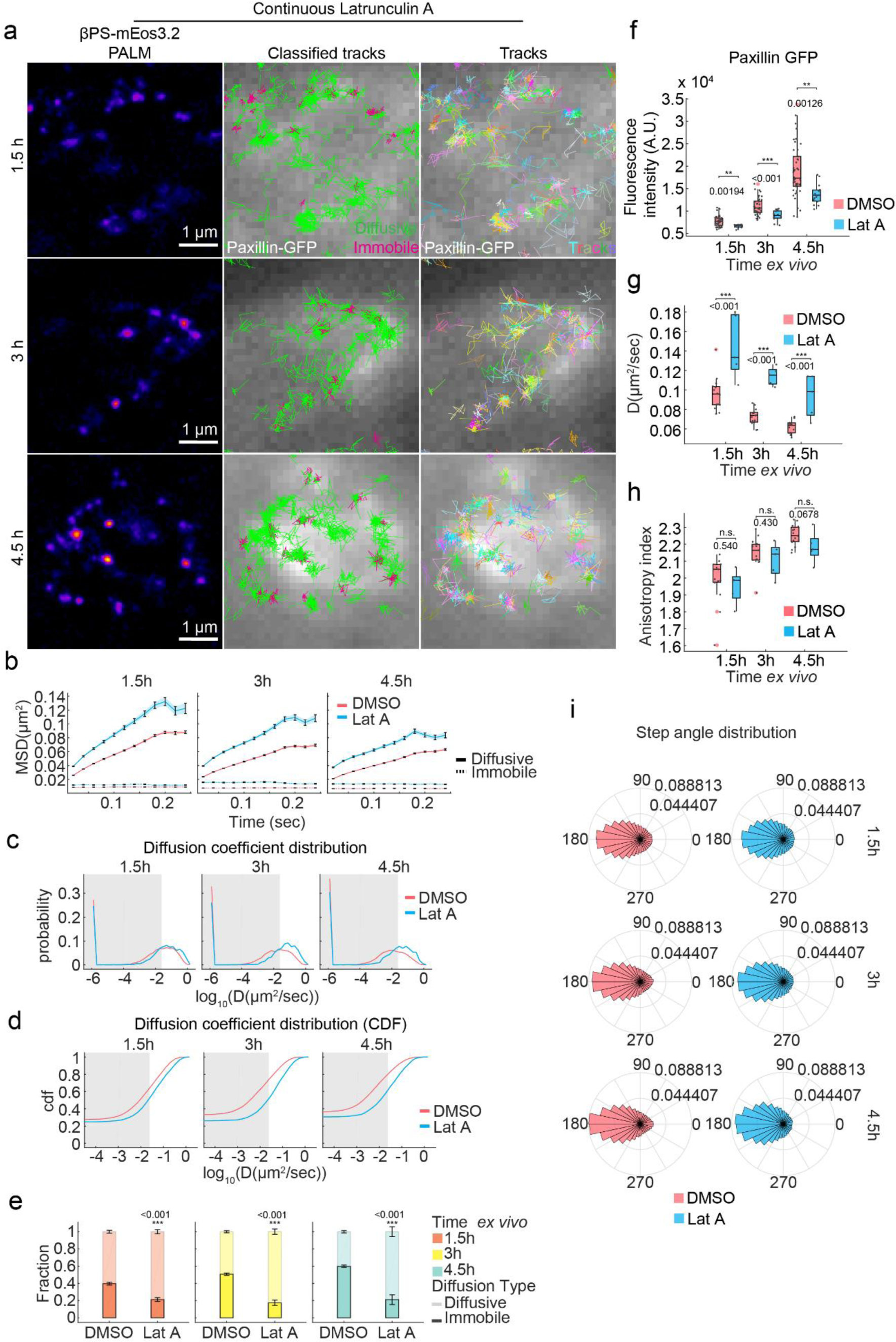
sptPALM tracking of βPS-mEos3.2 in developing adhesion with continuous Latrunculin A treatment. **a**, sptPALM tracking of βPS-mEos3.2 with continuous Latrunculin A (LatA) treatment from after dissection. Left column: reconstructed PALM images; centre column: tracks are characterized into diffusive (green) or immobile (red); Right column: tracks are plotted with a random color. Scale bars, 1 μm. **b**, Mean square displacement (MSD) of diffusive and immobile βPS -mEos3.2. **c**, Diffusion coefficient distribution of βPS -mEos3.2 tracks (probability). **d**, Diffusion coefficient distribution of βPS -mEos3.2 tracks (cdf). **e**, Fraction of Immobile and diffusive tracks. n = 15 embryos for DMSO 1.5h, 3h and 4.5h, n = 5 embryos for LatA 1.5h, 3h and 4.5h. **f**, Paxillin-GFP fluorescence intensity. n = 42 adhesions for DMSO 1.5h, n = 41 for DMSO 3h, n = 46 for DMSO 4.5h, n = 12 for LatA 1.5h, n = 14 for LatA 3h, n = 15 for LatA 4.5h. **g**, Diffusion coefficient n = 15 embryos for DMSO 1.5h, 3h and 4.5h, n = 5 embryos for LatA 1.5h, 3h and 4.5h. **h**, Step angle anisotropy. n = 15 embryos for DMSO 1.5h, 3h and 4.5h, n = 5 embryos for LatA 1.5h, 3h and 4.5h. **i**, Step angle distribution.

**Extended Data Figure 12.**
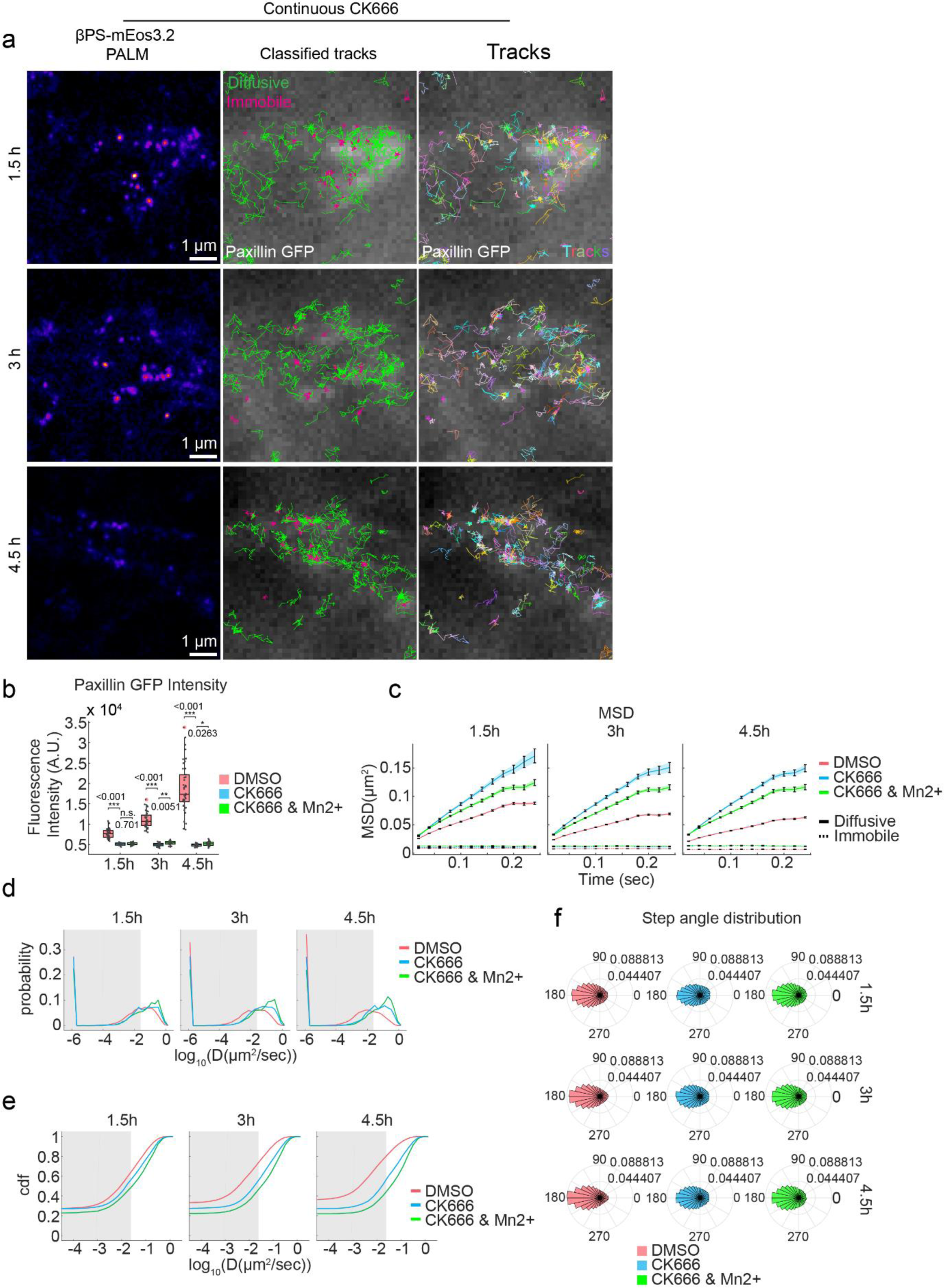
sptPALM tracking of βPS-mEos3.2 in developing adhesion with continuous CK666 treatment and Mn^2+^ stimulation. **a**, sptPALM tracking of βPS-mEos3.2 with continuous CK666 treatment from after dissection. Left column: reconstructed PALM images; centre column: tracks are characterized into diffusive (green) or immobile (red); Right column: tracks are plotted with a random colour. Scale bars, 1 μm. **b**, Paxillin-GFP fluorescence intensity. n = 42 adhesions for DMSO 1.5h, n = 41 for DMSO 3h, n = 46 for DMSO 4.5h, n = 18 for CK666 1.5h, n = 26 for CK666 3h, n = 21 for CK666 4.5h, n = 21 for CK666 & Mn^2+^ 1.5h, n = 20 for CK666 & Mn^2+^ 3h, n = 22 for CK666 & Mn^2+^ 4.5h. **c**, Mean square displacement (MSD) of diffusive and immobile βPS-mEos3.2. **d**, Diffusion coefficient distribution of βPS -mEos3.2 tracks (probability). **e**, Diffusion coefficient distribution of βPS -mEos3.2 tracks (cdf). **f**, Step angle distribution.

**Extended Data Figure 13.**
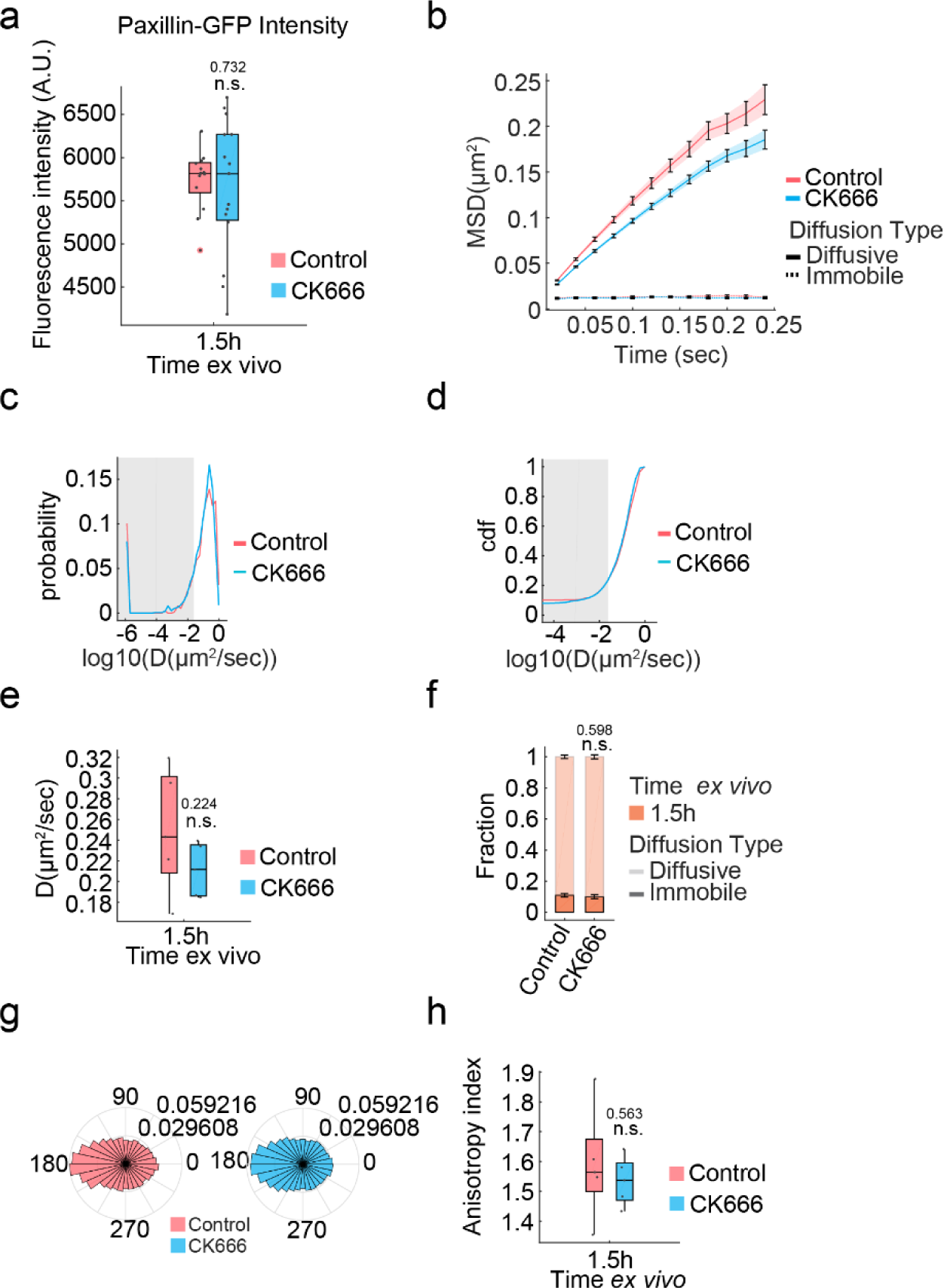
sptPALM tracking of βPS-mEos3.2 in cell body with CK666 treatment. **a**, Paxillin-GFP fluorescence intensity. n = 13 adhesions for control, n = 15 for CK666. **b**, Mean square displacement (MSD) of diffusive and immobile βPS-mEos3.2. **c**, Diffusion coefficient distribution of βPS -mEos3.2 tracks (probability). **d**, Diffusion coefficient distribution of βPS -mEos3.2 tracks (cdf). **e**, Diffusion coefficient. n = 5 embryos for control, n = 5 for CK666. **f**, Fraction of Immobile and diffusive tracks. n = 5 embryos for control, n = 5 for CK666 (mean ± s.e.m.). **g**, Step angle distribution. **h**, Step angle anisotropy. n = 5 embryos for control, n = 5 for CK666.

**Extended Data Figure 14.**
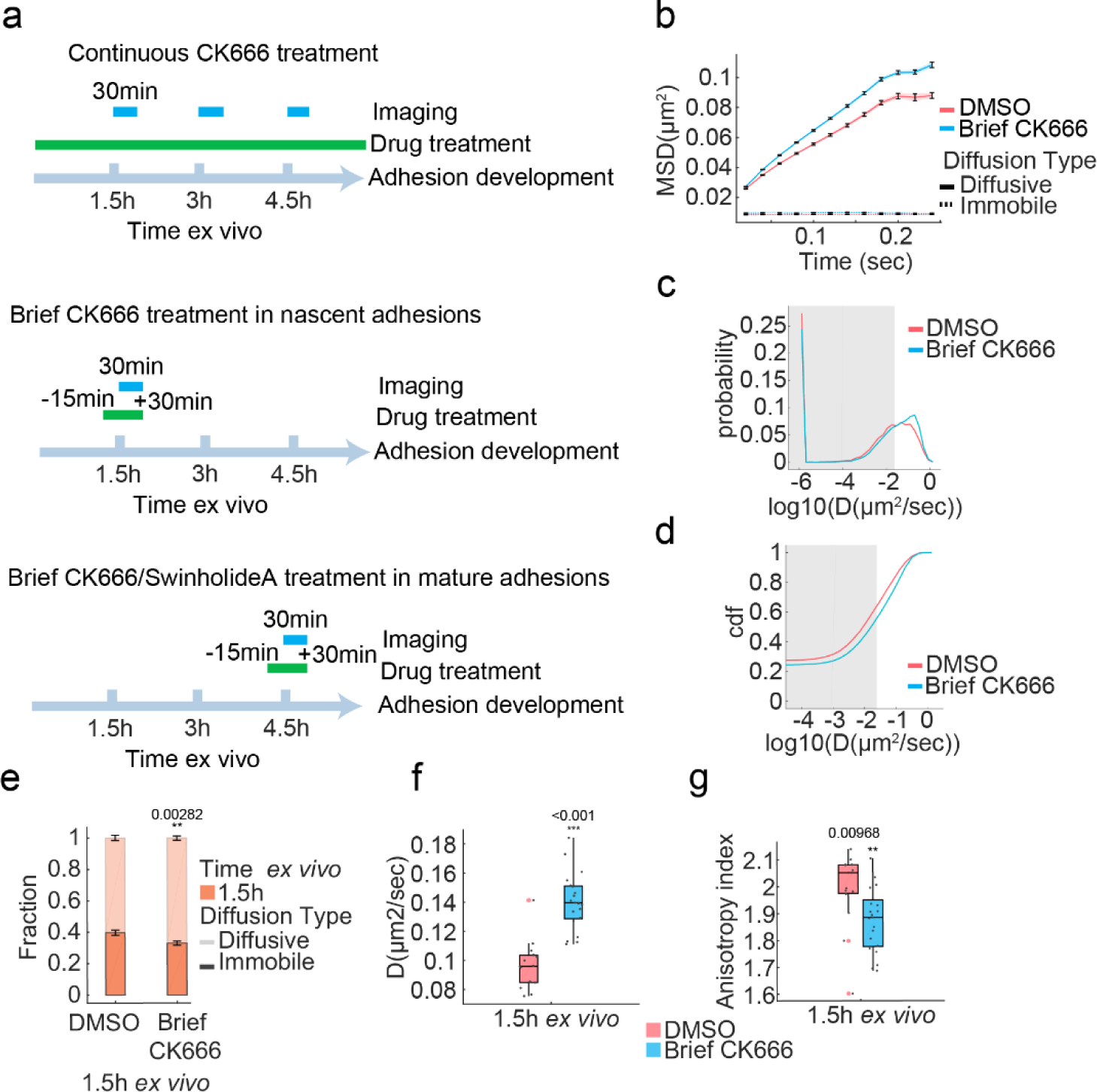
sptPALM tracking of βPS-mEos3.2 in 1.5h adhesions with brief CK666 treatment. **a**, Illustration showing experimental scheme for continuous and brief drug treatment and imaging during adhesion development. **b**, Mean square displacement (MSD) of diffusive and immobile βPS-mEos3.2. **c**, Diffusion coefficient distribution of βPS -mEos3.2 tracks (probability). **d**, Diffusion coefficient distribution of βPS -mEos3.2 tracks (cdf). **e**, Fraction of immobile and Diffusive trajectories for ꞵPS-mEos3.2 (mean ± s.e.m.). **f**, Diffusion coefficient of diffusive ꞵPS-mEos3.2 molecules. **g**, Step angle anisotropy for ꞵPS -mEos3.2.

**Extended Data Figure 15.**
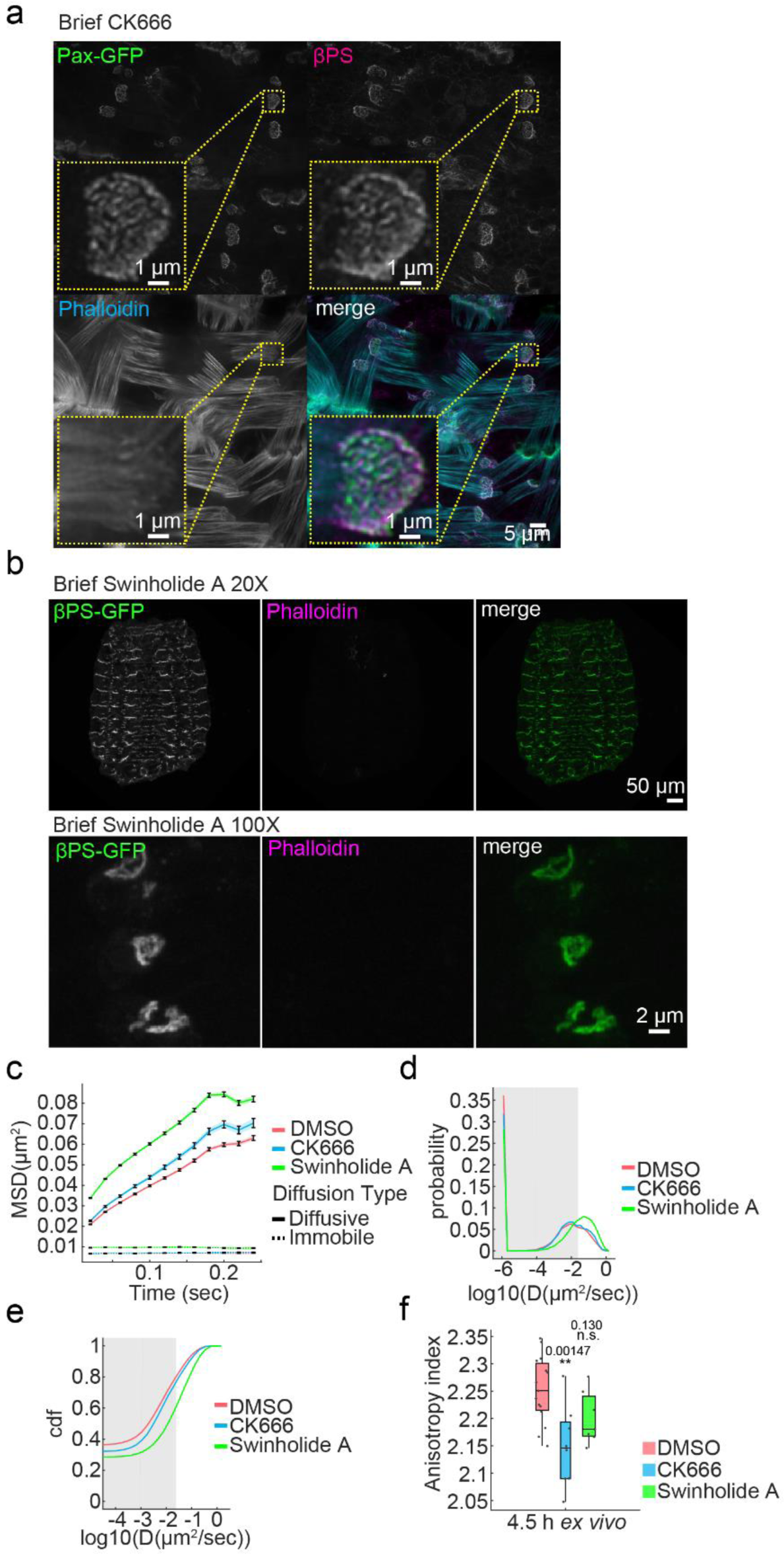
Brief CK666 and Swinholide A treatment in mature adhesions. **a**, Brief CK666 treatment in 4.5h *ex vivo* adhesions Green: Paxillin-GFP, magenta: βPS, cyan: phalloidin. Scalebar, 5 μm. Inset scalebars, 1 μm. **b**, Brief Swinholide A treatment in 4.5h ex vivo adhesions. Green: βPS-GFP, magenta: phalloidin. Scalebar, 50 μm for 20X, 2 μm for 100X; **c**, Mean square displacement (MSD) of diffusive and immobile βPS-mEos3.2. **d**, Diffusion coefficient distribution of βPS -mEos3.2 tracks (probability). **e**, Diffusion coefficient distribution of βPS -mEos3.2 tracks (cdf). **f**, Step angle anisotropy for ꞵPS - mEos3.2. n = 15 embryos for DMSO, n = 8 for CK666, n = 7 for Swinholide A.

**Extended Data Figure 16.**
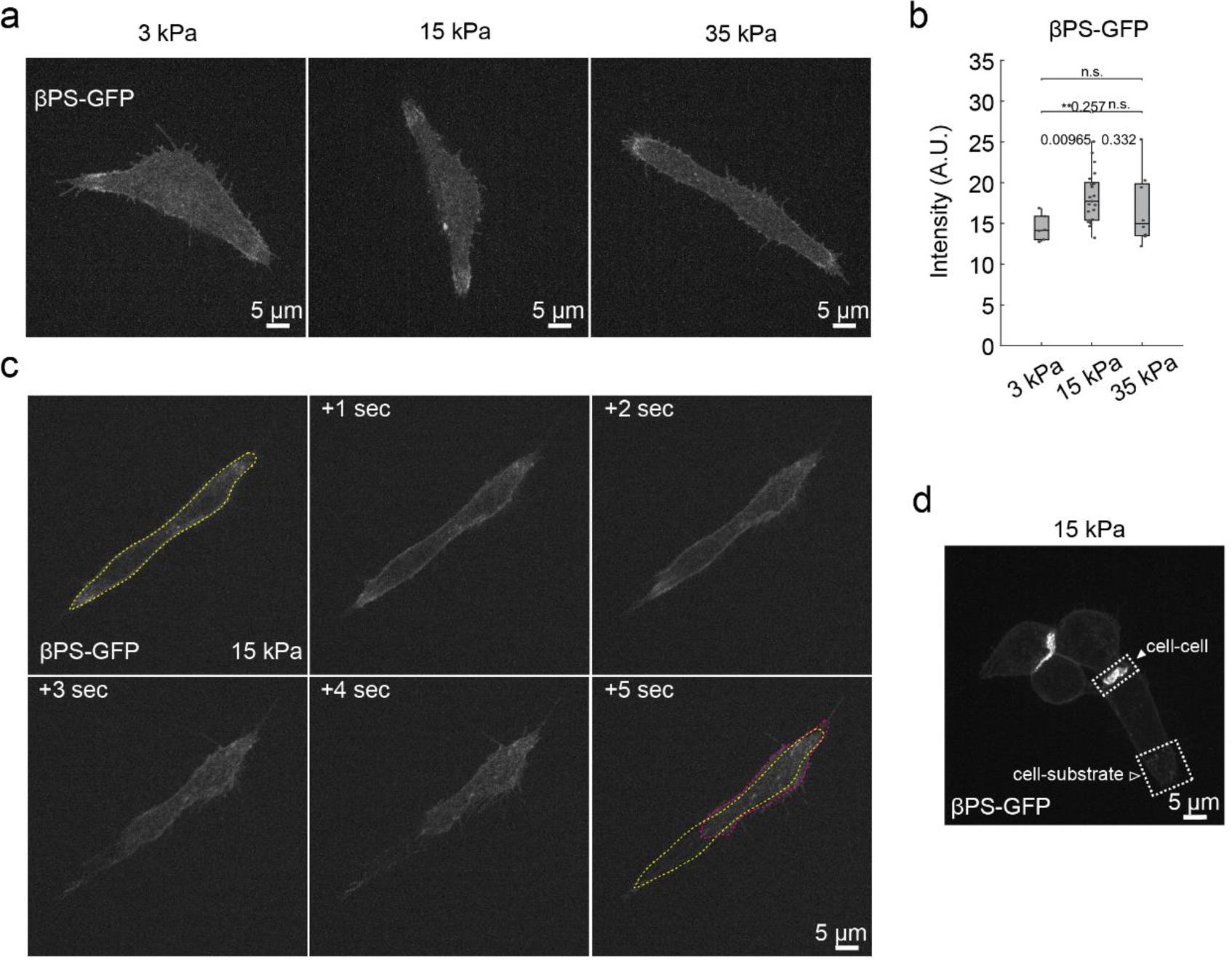
Isolated muscle cell adhesion development on PDMS gels of different rigidity. **a**, Isolated muscle cell and adhesion (βPS-GFP) morphology on PDMS gels of rigidities 3 kPa, 15 kPa and 35 kPa. Scalebar, 5 μm. **b**, βPS-GFP fluorescence intensity. n = 6 cells for 3kPa, n = 21 cells for 15kPa, n = 8 cells for 35kPa. **c**, Spontaneous muscle contraction leads to cell detachment on 15 kPa DMSO gel. Yellow dashed line: original outline of cell, magenta dashed line: cell outline after contraction Scalebar, 5 μm. **d**, Adhesion development between cells and between cell and substrate on 15 kPa PDMS gel. Scalebar, 5 μm.

**Extended Data Figure 17.**
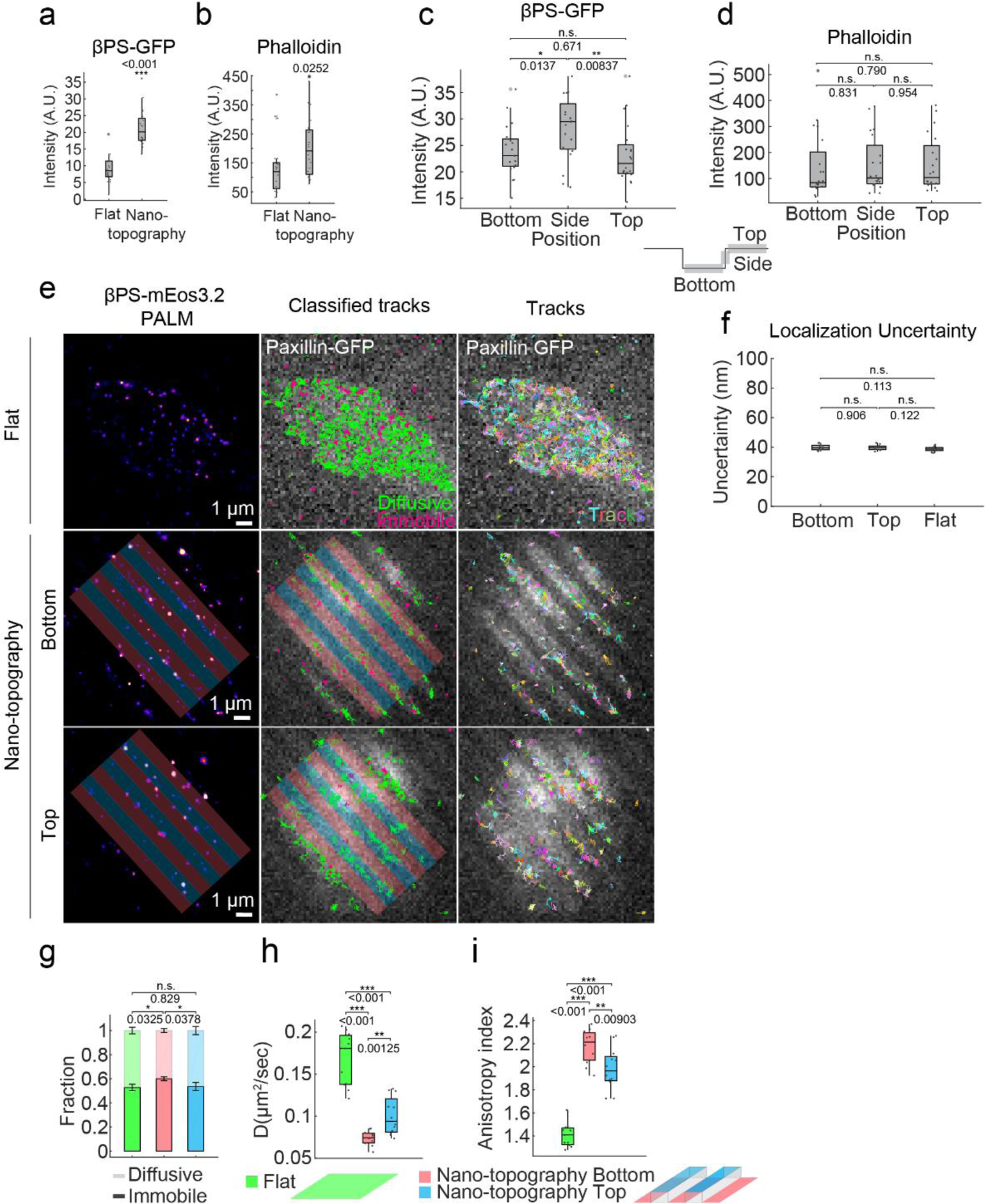
Nanotopography promotes adhesion formation and confinement of integrin diffusion. **a**, Fluorescence intensity of βPS-GFP in adhesions of isolated muscle cells on Curi Bio flat and nanotopography substrates. n = 21 cells for flat, n = 20 cells for nanotopography. **b**, Fluorescence intensity of phalloidin-Alexa647 in adhesions of isolated muscle cells on Curi Bio flat and nanotopography substrates. n = 21 cells for flat, n = 20 cells for nanotopography. **c**, Fluorescence intensity of βPS-GFP in adhesions of isolated muscle cells on Curi Bio nanotopography bottom, side and top regions. n = 20 cells for bottom, side and top. **d**, Fluorescence intensity of phalloidin-Alexa647 in adhesions of isolated muscle cells on Curi Bio nanotopography bottom, side and top regions. n = 20 cells for bottom, side and top. **e**, sptPALM tracking of βPS-mEos3.2 in isolated cells on flat, nanotopography bottom and top surfaces. Left column: reconstructed PALM images; centre column: tracks are characterized into diffusive (green) or immobile (magenta); Right column: tracks are plotted with a random colour. Bottom parts are shaded red and top parts are shaded blue. Scale bars, 1 μm. **f**, Localization uncertainty on flat, nanotopography bottom and top surfaces. n = 10 cells for bottom and top, n = 14 for flat. **g**, Fraction of Immobile and diffusive tracks. n = 11 embryos for flat, n = 11 for bottom, n = 12 for top. **h**, Diffusion coefficient. n = 11 embryos for flat, n = 11 for bottom, n = 12 for top. **i**, Step angle anisotropy. n = 11 embryos for flat, n = 11 for bottom, n = 12 for top.

**Extended Data Figure 18.**
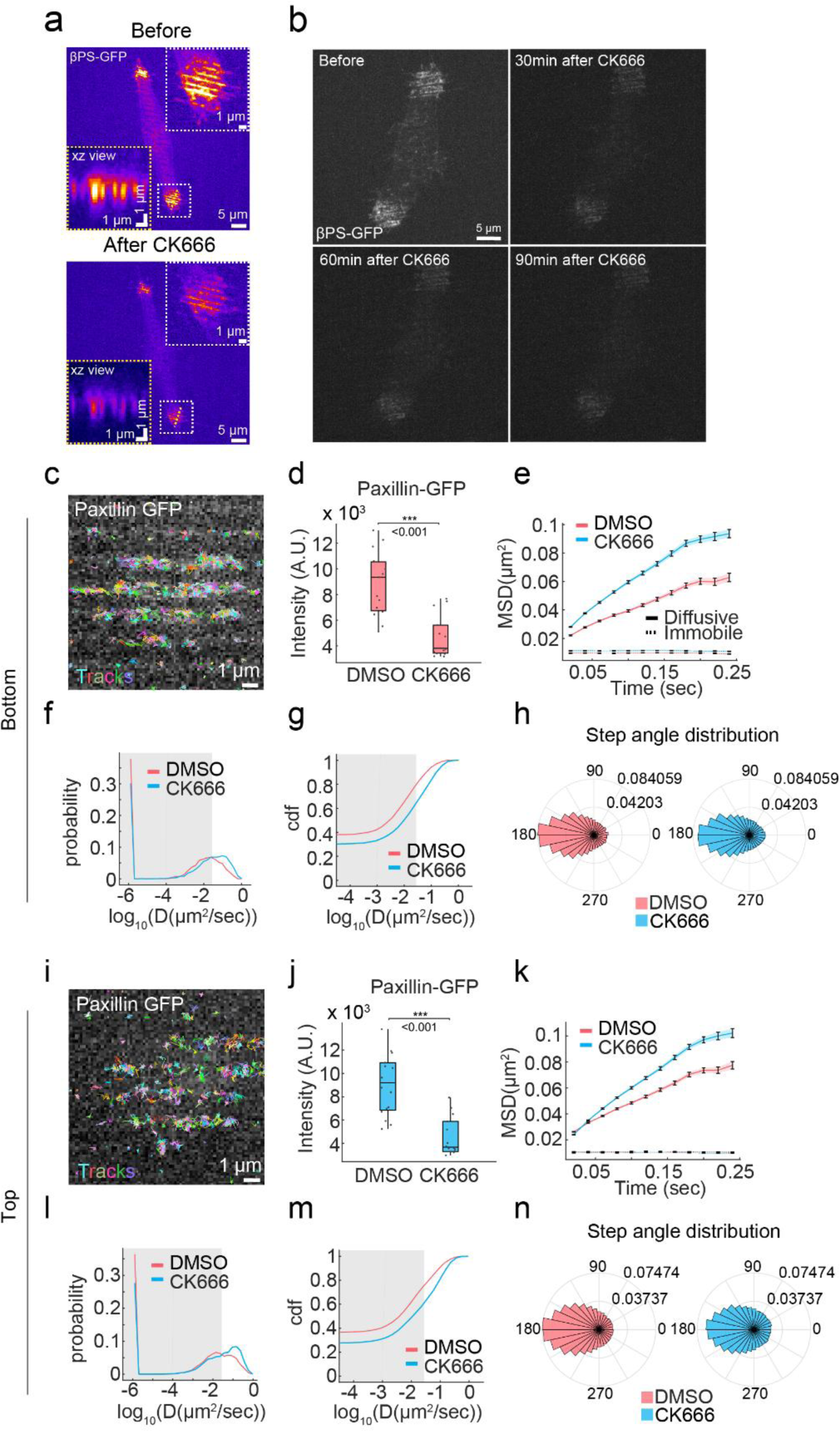
Nanotopography driven adhesion formation and confinement of integrin diffusion depends on Arp2/3 induced actin polymerization. **a**, βPS -GFP images of isolated muscle cells on nano-pattern before (top) and after (bottom) brief CK666 treatment (15 min). Scale bars, 5 μm. Inset scale bars, 1 μm. xz view scale bars (horizontal and vertical), 1 μm. **b**, βPS -GFP images of isolated muscle cells on nano-pattern before and after continuous CK666 treatment. Scale bars, 5 μm. **c**, sptPALM tracking of βPS-mEos3.2 in isolated cells on nanotopography bottom surface with continuous CK666 treatment. Tracks are plotted with a random colour. Scale bar, 1 μm. **d**, Paxillin-GFP fluorescence intensity. n = 15 cells for DMSO, n = 14 for CK666. **e**, Mean square displacement (MSD) of diffusive and immobile βPS-mEos3.2. **f**, Diffusion coefficient distribution of βPS -mEos3.2 tracks (probability). **g**, Diffusion coefficient distribution of βPS -mEos3.2 tracks (cdf). **h**, Step angle distribution. **i**, sptPALM tracking of βPS-mEos3.2 in isolated cells on nanotopography top surface with continuous CK666 treatment. Tracks are plotted with a random colour. Scale bar, 1 μm. **j**, Paxillin-GFP fluorescence intensity. n = 16 cells for DMSO, n = 14 for CK666. **k**, Mean square displacement (MSD) of diffusive and immobile βPS-mEos3.2. **l**, Diffusion coefficient distribution of βPS -mEos3.2 tracks (probability). **m**, Diffusion coefficient distribution of βPS -mEos3.2 tracks (cdf). **n**, Step angle distribution.

**Extended Data Figure 19.**
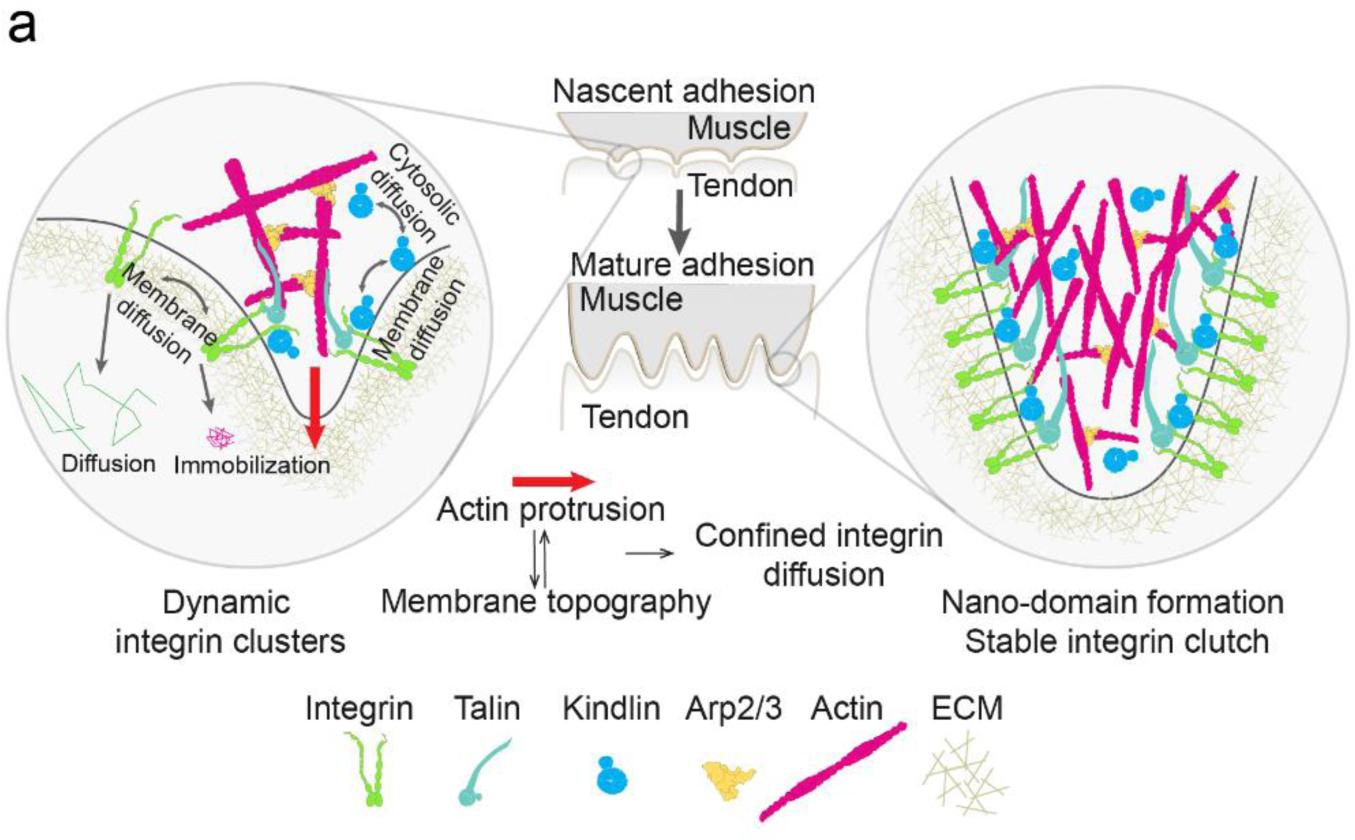
Conceptual model. **a**, Illustration showing conceptual model of actin dynamics and regulation of integrin diffusion by nanotopography during adhesion development.

### Supplementary Movies

**Supplementary Movie 1.**

Development of muscle attachment site (MAS) in an embryo fillet over 6 hours ex vivo with paxillin-GFP labelling, 20X objective, spinning disk confocal, 10 min per frame. Scale bar, 50 µm.

**Supplementary Movie 2.**

Development of MAS over 8 hours *ex vivo* with paxillin-GFP labelling, 100X objective, spinning disk confocal, 10 min per frame. Scale bar, 2 µm.

**Supplementary Movie 3.**

20X DIC movie showing spontaneous muscle contraction in an embryo fillet (>6h *ex vivo*), 3X real time. Scale bar, 50 µm.

**Supplementary Movie 4.**

Movie showing the spontaneous contraction of individual muscles in an embryo fillet labelled with utrophin-GFP (>6h *ex vivo*), 20X objective, spinning disk confocal, 3X real time. Scale bar, 50 µm.

**Supplementary Movie 5.**

3D reconstruction from Modloc 3D super-resolution imaging of integrin clusters from adhesions at 0.5h, 1.5h, and 4.5h *ex vivo*. See Fig. 1d for scale.

**Supplementary Movie 6.**

Continuous CK666 treatment disassembles mature adhesions (βPS-GFP) and abolishes MAS development, 100X objective, spinning disk confocal. Scale bar, 2 µm.

**Supplementary Movie 7.**

3D reconstruction of Utrophin-GFP labelled developing MAS shows dynamic actin protrusions in early adhesions and stable topographical structures in mature adhesions, 100X objective, spinning disk confocal. Scale bar, 5 µm.

**Supplementary Movie 8.**

Integrin foci move and fuse with existing structures in developing MAS, 100X objective, spinning disk confocal. Scale bar, 2 µm.

**Supplementary Movie 9.**

Actin (Utrophin-GFP) puncta dynamics in developing MAS at different stages, 100X objective, spinning disk confocal. Scale bars, 1 µm.

**Supplementary Movie 10.**

CK666 treatment abolishes actin (Utrophin-GFP) puncta dynamics in developing MAS (1.5h *ex vivo*), 100X objective, spinning disk confocal. Scale bar, 1 µm.

**Supplementary Movie 11.**

CK666 treatment abolishes integrin (βPS-GFP) foci dynamics in developing MAS (1.5h *ex vivo*), 100X objective, spinning disk confocal. Scale bar, 1 µm.

**Supplementary Movie 12.**

Example showing sptPALM imaging of integrin (βPS-mEos3.2, magenta, sptPALM Hilo illumination) molecule diffusion and immobilization in developing adhesion (paxillin-GFP, green, widefield) at 1.5h, 3h and 4.5h *ex vivo*, 100X objective. Scale bar, 2 µm.

**Supplementary Movie 13.**

Brief treatment of Swinholide A disrupts mature adhesion (βPS-mEos3.2, 4.5h *ex vivo*) morphology, 100X objective, spinning disk confocal. Movie starts immediately after addition of Swinholide A. Scale bar, 2 µm.

**Supplementary Movie 14.**

Spontaneous contraction of isolated muscle cell on 15kPa PDMS gel causes detachment of immature adhesions (βPS-mEos3.2), 100X objective, spinning disk confocal. Scale bar, 5 µm.

